# Interconnected axes of phenotypic plasticity drive coordinated cellular behaviour and worse clinical outcomes in breast cancer

**DOI:** 10.1101/2025.11.23.688595

**Authors:** Ritesh Kumar Meena, Yijia Fan, Soundharya Ramu, Anjaney J. Pandey, Abhinav Kannan, Yashvita Subramanian, Yukta Subramanian, Jason T. George, Mohit Kumar Jolly

## Abstract

Phenotypic plasticity plays a key role in cancer progression and metastasis, enabling cancer cells to adapt and evolve, but precisely how distinct axes governing phenotypic plasticity interact to shape tumour progression and patient outcomes remains unclear. We investigated five major interconnected axes of plasticity in ER-positive (ER+) breast cancer: Metabolic Reprogramming, Epithelial-to-Mesenchymal Plasticity (EMP), Luminal–Basal (Lineage) Switching, Stemness, and Drug-resistance using network dynamics simulations, integrative bulk and single-cell transcriptomic analyses and patient survival analyses. We show that these axes are not independent but drive one another, forming two mutually inhibiting ‘teams’ of nodes enabling specified cellular behaviour. One team (favouring high glycolysis, stem-like, basal-like, mesenchymal/hybrid and tamoxifen-resistant phenotype) was found to be associated with aggressive progression and worse survival. On the other hand, the opposing team (favouring high oxidative phosphorylation, non-stem-like, luminal-like, epithelial and tamoxifen-sensitive phenotype) correlated with better outcomes. Importantly, altering one axis of plasticity often drove coordinated responses along other axes and vice versa. Our findings establish phenotypic plasticity in cancer as a coordinated, multi-axis dynamical process, thus suggesting novel strategies to disrupt systems-level reprogramming enabling metastasis and therapeutic resistance.

## 1 Introduction

Cancer is a group of diseases characterised by uncontrolled cell growth. A subset of cancer cells have the potential to detach from the primary tumor, enter the blood or lymphatic system, and establish secondary tumors in distant organs through a process called metastasis. Although 99.9% of these disseminated cells fail to metastasise [1], the remaining 0.1% that do account for the majority of cancer-related mortality, thus making metastasis a major unsolved clinical challenge. Furthermore, phenotypic plasticity — the ability of cancer cells to reversibly alter their phenotype without altering their genomic architecture — is one of the major drivers of metastasis [2].

Arguably, the most extensively studied form of phenotypic plasticity involves an Epithelial-to-Mesenchymal Transition (EMT) and its reverse Mesenchymal-to-Epithelial Transition (MET). During EMT, cells at least partially lose epithelial features such as cell-cell adhesion and acquire mesenchymal traits, including increased motility and invasiveness. EMT and MET are both more commonly now referred to as Epithelial-to-Mesenchymal Plasticity (EMP) [3]. Other axes of phenotypic plasticity relevant to the cancer context include reversible transitions between cancer stem cells & non-cancer stem cells [4], metabolic reprogramming between Oxphos & Glycolysis [5] and lineage plasticity as observed, for instance, in ER+ breast cancer and bladder cancer, from luminal to basal [6].

Recent experimental and theoretical studies have identified molecules such as HIF1*α* and AMPK, as key regulators of metabolic plasticity, to be differentially expressed during EMT [7, 8, 9]; HIF1*α* is typically upregulated [10, 11], whereas AMPK is downregulated [12]. HIF1*α* knock-down can abrogate EMT and consequently reduce metastasis in a mouse model [13]. Conversely, knockdown of ZEB1, an EMT-inducing transcription factor, can alter the metabolic composition of cells and impact ferroptosis sensitivity [14]. Furthermore, in breast cancer, both hybrid E/M and mesenchymal phenotypes are associated with enhanced resistance to estrogen therapy and increased immunosuppression through mechanisms such as PD-L1 upregulation [15, 16, 17]. Together, these examples highlight the possibility of a potential crosstalk between drivers of interconnect axes of phenotypic plasticity that influence one another. While the dynamics of dual-axes associations such as EMT–drug resistance [18, 19], EMT–Stemness [20, 21], EMT–immune suppression [15, 22, 23], and EMT–metabolism [5, 24, 25] have been reported, studies addressing multiple axes (*>* 2) simultaneously remain limited. Given the pivotal role of phenotypic plasticity in driving metastasis, investigating this interconnected network of biological processes is of crucial importance and can aid in developing new combination therapies to more effectively target cancer cells.

In this study, we aim to understand the coordination between molecules regulating five different axes of phenotypic plasticity. We constructed a minimal Gene Regulatory Network (GRN) by identifying modules consisting of key genes associated with five different axes of phenotypic plasticity namely, Metabolic Reprogramming (related to the alteration of cellular metabolism to meet the high energetic and biosynthetic demands of rapidly proliferating tumour cells), Epithelial-to-Mesenchymal plasticity (related to the ability of cancer cells—to transition between epithelial and mesenchymal states), Stemness (related to reversible interconversion between cancer stem cells (CSCs) and non-stem cancer cells (non-CSCs)), Drug Resistance (related to dynamic and reversible transitions between drug-sensitive and drug-resistant cellular states) and specific to breast cancer Luminal–Basal (Lineage) switching (related to phenotypic plasticity in epithelial cancers where tumor cells can switch between luminal and basal lineage states). The proposed gene regulatory network was simulated to study the emergent phenotypic states due to the intra– and-module interactions. The simulations revealed the existence of two mutually repressing ‘teams’, one consisting of genes that promote EMT along with high-glycolysis, stem-like, drug-resistant, basal-like phenotype, whereas the other consisted of genes that have an inhibitory effect on EMT along with genes that promote a high-Oxphos, non-stem-like, drug-sensitive and luminal-like phenotype. Subsequently, GSEA performed on transcriptomic data showed that the induction of EMT or drug resistance not only altered the activity of EMT and drug response gene sets, but the effects of these alterations propagated across other axes of plasticity, including metabolism and luminal-basal programs. Furthermore, survival analysis revealed that these associations between different axes of phenotypic plasticity have implications in terms of therapeutic outcomes, wherein the coordinated activation of multiple axes together but not necessarily individual axes alone drives worse outcomes.

## 2 Results

### 2.1 Integrated gene regulatory network dynamics shows multi-modal distribution of phenotypic scores

The first step in understanding how different axes of cellular plasticity in cancer cells interact with one another was the identification of a minimal Gene Regulatory Network (GRN) which could explain experimentally observed results. Based on literature and prior work, a GRN was identified, where genes were represented as nodes grouped into five modules, namely: (1) EMT, (2) Stemness, (3) Tamoxifen-resistance, (4) Luminal-Basal Identity, and (5) Metabolic Reprogramming. This final network was composed of a total of 24 nodes and 106 edges (Figure 1a)[Supplementary Table 1].

**Figure 1:**
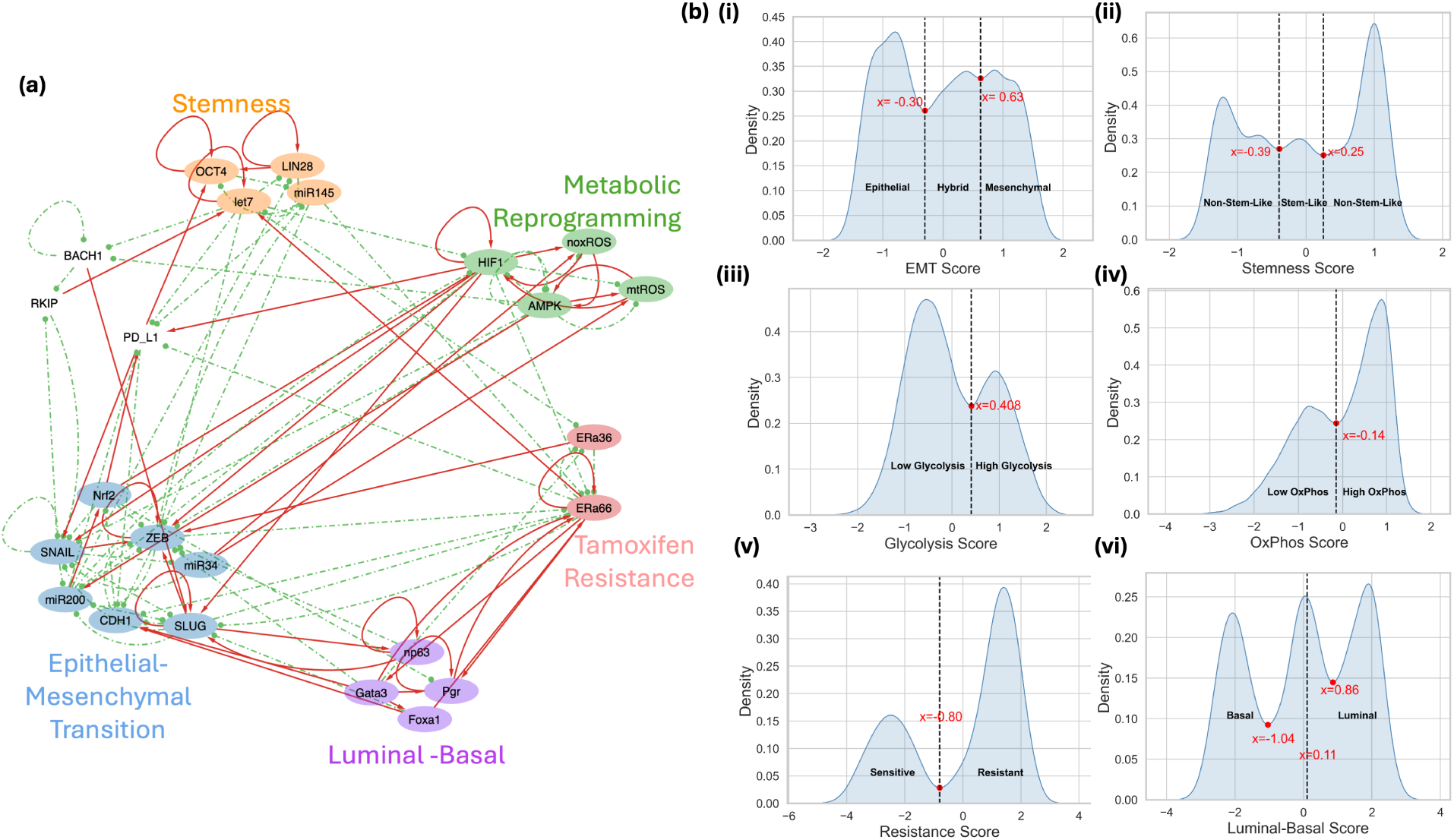
Gene regulatory network analysis for coupled axes of plasticity. (a) GRN showing interactions between nodes from different modules. Nodes in the network represent genes, and the edges between nodes represent regulations between respective nodes. Red arrows with pointed ends indicate activation; Green arrows with blunt ends indicate inhibition. (b)(i) Kernel density plot for EMT score (= (ZEB + SLUG – miR200 – CDH1)/4). (ii) Kernel density plot for Stemness score (= (OCT4 + LIN28 – mir145 – let7)/2). (iii) Kernel density plot for Glycolysis Score (= (noxROS + HIF1)/2). (iv) Kernel density plot for Oxphos score (= (AMPK + mtROS)/2). (v) Kernel density plot for Resistance Score (= (ERa36 – ERa66)/2). (vi) Kernel density plot for Luminal-Basal Score (= (ERα66 + GATA3 + PGR + FOXA1)/4–(NP63 + SLUG)/2)). Black dashed lines indicate the x-coordinate(s) of minima from the fitted Gaussians used for segregation of phenotypes for respective scores.

These five different modules consists of: **(1) The EMT-module**, included nodes that stabilize and promote the Mesenchymal phenotype (ZEB1, SNAI1/ SNAIL, SNAI2/ SLUG), the Epithelial phenotype (CDH1, miR200, miR34) and the Hybrid phenotype of cancer cells (NRF2)[26]; **(2) The Stemness module**, included stemness-promoting nodes (LIN-28 and OCT-7) and stemness-suppressing nodes (Let-7 and miR145), with the pairs LIN28–let-7 and miR-145–OCT4 engaged in mutual inhibitory loop [21, 27]; **(3) The Tamoxifen-resistance module**, included ER*α*66 and ER*α*36, two variants of estrogen receptor (ESR1). Higher levels of ER*α*66 are associated with tamoxifen-sensitive cell state, while that of ER*α*36 are associated with tamoxifen-resistant cell state [28, 19]; **(4) The Luminal–Basal module**, included nodes promoting a Luminal phenotype (ER*α*66, PGR, GATA3, and FOXA1) and nodes promoting a Basal phenotype (SLUG and np63) [29]; **(5)The Metabolic Reprogramming module**, included nodes related to Glycolysis (noxROS and HIF1*α*) and Oxidative Phosphorylation (AMPK and mtROS) [25]; In addition, the network contained three more nodes namely PD-L1, representing a central immune checkpoint molecule regulating immune escape of cancer cells, and toggle switch couple RKIP–BACH1 known to have roles in inhibition and promotion of EMT respectively [30, 31]. We note that the provided network is not meant to be exhaustive in terms of interaction between nodes, but it demonstrates a core network structure that may be sufficient to explain the correlation-based observations observed in the literature.

Next, to study the emergent properties of this GRN, we simulated it using RACIPE, a computational framework that solves a set of coupled ordinary differential equations (ODEs) [32], to examine the various phenotypic states enabled by the GRN by sampling an ensemble of kinetic parameter sets from a biologically relevant parameter range. (See RACIPE: Material and Methods).

Further, to obtain a more quantitative understanding of how interactions between different modules affected the population distribution in the simulation cohort, module-specific phenotypic scores were defined. The EMT score was defined as (ZEB+SLUG *−* miR200 *−* CDH1)*/*4; the Stemness (SN) score as (OCT4 +LIN28 *−* miR145 *−* let7)*/*2; the Glycolysis Score as (noxROS+HIF1)*/*2; the Oxphos score as (AMPK + mtROS)*/*2; the Luminal–Basal Score as (ER*α*66 + GATA3 + PGR + FOXA1)*/*4 *−* (NP63 + SLUG)*/*2; and the Resistance Score as (ER*α*36 *−* ER*α*66)*/*2. Inspection of the density distributions of these scores revealed patterns consistent with those reported in the literature for smaller and less complex networks. Specifically, the EMT score exhibited a trimodal distribution, corresponding to three distinct subpopulations (Figure 1b(i)): low scores (epithelial), intermediate scores (hybrid E/M), and high scores (mesenchymal), similar to the distributions obtained earlier [30]. Likewise, cells with extremely low or extremely high LIN28 and correspondingly OCT4 levels are associated with a non-stem-like phenotype, whereas those with intermediate levels of OCT4 correspond to a stem-like phenotype (Figure 1b(ii)) [33], again aligning with results obtained in earlier studies [30].

The density distributions for Glycolysis and Oxphos scores (Figure 1b(iii), Figure 1b(iv)) were predominantly bimodal, resulting in cell categorisation into high/low Glycolysis and Oxphos. Because cells with high ER*α*36 and low ER*α*66 expression are known to exhibit greater resistance to Tamoxifen[28], the bimodal distribution of the Resistance Score was used to classify cells with high scores as Resistant and those with low scores as Sensitive (Figure 1b(v)), similiar to previous stud-ies [30, 19]. For Luminal–Basal Score, a trimodal distribution was observed; hence the median was used to label cells as Luminal (with high Luminal–Basal Score) and Basal (with low Luminal–Basal Score), as reported previously [29] (Figure 1b(vi)).

These observations of the bimodal density distributions of individual scores, which remain unchanged upon integrating multiple modules into a single network, suggest that these different axes of phenotypic plasticity operate as components of a larger and coordinated system rather than independent entities.

### 2.2 Opposing ‘Teams’ of genes define distinct phenotypic states associated with different axes of phenotypic plasticity

Before analysing the simulation results, we inspected the outcomes of regulations between nodes within and across different modules in the GRN. For this purpose, two matrices were constructed: a) Adjacency matrix (Figure 2a(i)), representing the direct interactions from source nodes (rows) to target nodes (columns) and b) Influence matrix, which quantifies the cumulative effect of direct and indirect interactions between each pair of nodes. Both matrices were derived solely from the network topology without doing any simulations (see Adjacency matrix and Influence matrix, Materials and Methods). Teams’ analysis on the Influence matrix (Figure 2a(ii)) revealed the existence of two teams of molecular players which effectively activate members of the same team and inhibit members of the opposing team (see Teams’ Analysis: Material and Methods). The first team consisted of pro-hybrid/ mesenchymal nodes like ZEB1, SNAIL, NRF2 and SLUG, along with molecules that are associated with a basal-like phenotype (np63 and SLUG), high-glycolysis (noxROS and HIF1), stem-like features (OCT4 and LIN28), an immune regulator (PD-L1) and a drug-resistant phenotype (ERa36). The second team consisted of pro-epithelial molecules, such as miR200 and CDH1, along with molecules that correspond to a luminal-like phenotype (GATA3, FOXA1, Pgr, and miR145), high-Oxphos (AMPK and mtROS), non-stem-like features (miR145 and let-7), and a drug-sensitive phenotype (ERa66).

**Figure 2:**
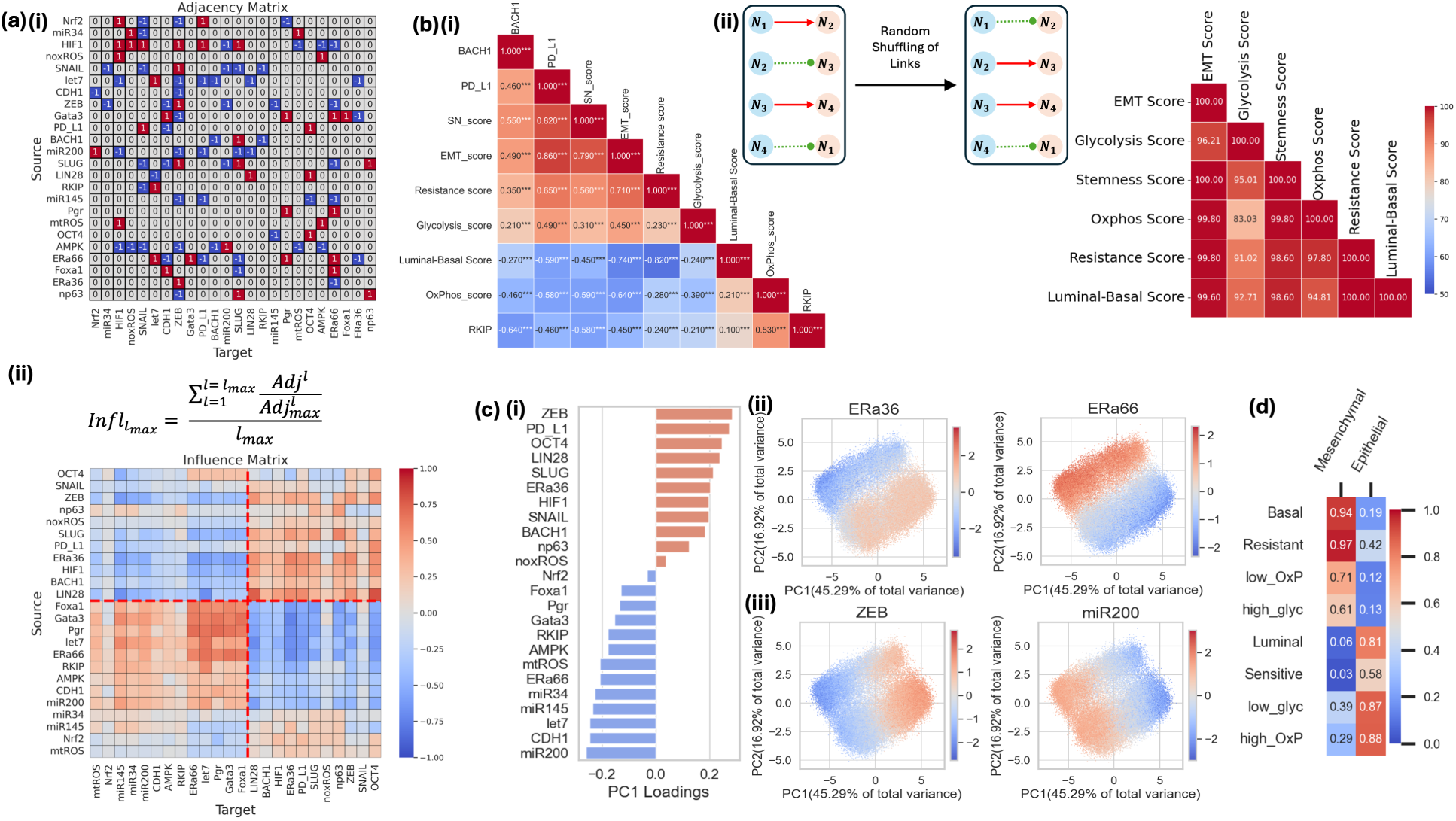
Emergent associations between modules as an outcome of Interactions between nodes from different modules. (a)(i) Adjacency matrix for the network shown in Figure 1(a). (ii) Influence matrix using path length = 10 for the network shown in Figure 1(a). (b)(i) Correlation matrix between different scores and Gene expression levels. (ii) Network randomisation schematic and heatmap showing percentile rank of the WT network in the cohort of 500 random networks. (c) Results from principal component analysis applied to simulations (i) Vertical bar plot showing the PC1 loading coefficients of different genes in the network. (ii, iii) Scatterplots of PC1 (capturing 45.29% variance) vs PC2 (capturing 16.92% variance) shaded by z-normalised score values for genes associated with Drug-Resistance module (ii) and EMT module (iii). (d) Heatmap for the conditional probability of phenotypes associated with different axes of plasticity, given the epithelial and mesenchymal phenotypes of the cell.

Following the z-normalisation of steady-state solutions obtained by RACIPE simulation results, we generated a pairwise correlation analysis among the defined scores and the expression values of RKIP, BACH1, and PD-L1. Hierarchical clustering of the matrix revealed two clusters (Figure 2b(i)). The SN-score, Resistance score, and Glycolysis score clustered with the EMT-score, forming one group. In contrast, the Luminal–Basal score and Oxphos score clustered together as a separate group. Scores within both clusters showed a positive correlation with scores in the same group and a negative correlation with scores in the other group. The expression levels of PD-L1, a surface receptor molecule known to enable immune evasion, clustered with EMT, SN, Resistance, and Glycolysis scores. Furthermore, expression of BACH1, a pro-EMT transcription factor, clustered with the EMT group. In contrast, RKIP (Raf Kinase Inhibitor Protein) expression clustered with the Oxphos and Luminal–Basal group. These observations highlight that cellular programmes associated with different axes of cellular plasticity considered here formed two groups, one group works in unison with EMT and another in opposition to it. These results were in complete agreement with results from the ‘Teams’ analysis of the GRN.

To assess whether the presence of these ‘teams’ and clustering of associated phenotypic sores was unique to the particular wild-type (WT) network we considered, we generated 500 random networks by randomly shuffling edges among the nodes (see Generation of Random Networks: Materials and Methods). Comparison of network team-strength further revealed that the WT network exhibited substantially higher team strength than any of the 500 random networks (Figure S1a), suggesting that the presence of teams-behavior is a property specific to the WT network considered here. Next, we simulated each random network using RACIPE, and performed pairwise Spearman correlation analyses for steady-state solutions obtained for each network, as previously done for the WT network. To quantify the relative robustness of the WT network, we conducted a comparative percentile analysis across different score-pairs of the WT network with the random networks (Figure 2b(ii), Figure S1b (i)-(xv)). For 14 out of 15 combination pairs, the percentile rank of the WT correlation coefficient exceeded 90%. Importantly, it was observed that no single random network displayed stronger correlations than the WT network across all scores. These results strongly support that the organisation of molecular players into ‘teams’ is not a random occurrence but rather a distinctive feature of the WT-network topology that promotes coordination among these different axes of phenotypic plasticity.

Principal component analysis (PCA) was performed on the z-normalised RACIPE solutions. Over 60% of the variance could be captured by the first two principal components (PCs): PC1 accounted for 45.29% of the variance, while PC2 accounted for 16.92%. The loading coefficients of each node in the network along the PC1 axis were plotted, which again revealed two groups: one with all positive loading coefficients, and the other with negative ones (Figure 2c). Intriguingly, the composition of these two groups was exactly the same as that of the ones obtained from the teams’ analysis of the influence matrix. The cluster with pro-mesenchymal nodes/genes was found to have a positive PC1 coefficient, whereas the cluster with pro-epithelial nodes had a negative PC1 coefficient. Moreover, nodes with a strong biological association with phenotype represented by Team-1, such as ZEB1 (mesenchymal), PD-L1 (immune-evasive), OCT4 and LIN28 (stem-like), were found to have highly positive PC1 loading coefficients. In contrast, nodes with a strong biological association with an epithelial phenotype, such as CDH1, miR200, miR145, let7, and miR34, showed highly negative PC1 loading coefficients. Interestingly, NRF2, a well-known promoter of the hybrid phenotype in cancer cells, was observed to have a PC1 coefficient of very small magnitude. Next, we projected the gene expression profiles from the simulation cohort onto the PC1–PC2 space to examine the module-specific expression patterns of genes. Comparative analysis revealed the existence of an antagonistic relationship between genes across the ‘teams’, largely irrespective of the biological module considered. For the Tamoxifen-resistance module, ER*α*36 and ER*α*66 exhibited largely exclusive expression (Figure 2c(ii)). A similar antagonism was observed in EMT genes (ZEB vs. miR200) (Figure 2c(iii)), Stemness module genes (pro-stemness LIN28/OCT4 vs. stemness-suppressing let7/miR145), Luminal-Basal module genes (Gata3/Pgr vs. SLUG/np63) and Metabolic-Reprogramming genes (mtROS vs. HIF1). RKIP and BACH1 also show a comparable antagonism, where cell populations with high RKIP expression displayed low BACH1 levels, while populations with high BACH1 expression showed low RKIP levels (Figure S2b(i-iv)).

In addition to overlapping expression patterns, these plots also highlight synchronous expression patterns in genes belonging to different phenotypic plasticity modules. The expression of the tamoxifen-resistance gene (Era36) overlapped with basal genes (SLUG/ np63), while the expression of the tamoxifen-sensitive gene (Era66) overlapped with luminal genes (Gata3/ Pgr). Similar overlaps were observed where pro-EMT genes (ZEB1) overlapped with pro-stemness (LIN28/OCT4) and pro-glycolysis/ hypoxia genes (HIF1) while pro-epithelial genes(miR200), overlapped with stemness-suppressing (Gata3/ Pgr) and pro-Oxphos genes (mtROS). Together, these results highlighted the existence of coordinated regulation in gene expression across all five axes of phenotypic plasticity. Notably, we observe a subpopulation of cells along the EMT/ Tamoxifen-resistance axis and the EMT/ Luminal-Basal axis with mixed behaviour, suggesting the presence of mixed subpopulations reflecting heterogeneity in population structure.

Furthermore, to better understand how different emergent phenotypes coexist with EMT, a conditional probability analysis was conducted to quantify the probability of a certain phenotype, given a cell being mesenchymal or epithelial. It was observed that mesenchymal cells had a higher probability of showing a basal, tamoxifen-resistant, low-Oxphos and high-glycolysis phenotype (Figure 2d). In contrast, epithelial cells had a higher probability of exhibiting a luminal, tamoxifen-sensitive, high-Oxphos and low-glycolysis phenotype, i.e. completely opposite to the characteristics of mesenchymal cells.

Collectively, these results indicate that there are distinct cellular programmes, such as glycolysis, oxidative phosphorylation and drug sensitivity, that exhibit associations with one another and with EMT, functioning either in synchrony or in opposition. This reflects the well-established intra-tumoral heterogeneity in cancer cell populations, which may possibly be a direct emergent property of GRN dealing with multiple axes of phenotypic plasticity.

### 2.3 Bulk transcriptomic analysis suggests coordinated transitions across axes of plasticity

We next asked the question whether these co-expression patterns predicted by mathematical modelling of underlying GRNs are observed in transcriptomic data for cell lines and/or patient samples, and whether changes triggered along one axis can perturb the other, for instance, can EMT induction lead to tamoxifen resistance and/or vice-versa.

First, we investigated the RNA-seq data of MCF7 (an ER+ breast cancer cell line) cells in which the estrogen receptor (ER) has been silenced, GSE27473 [34]. Reduced expression of ESR1– gene that codes for estrogen receptor– is an established determinant of tamoxifen resistance in ER+ breast cancer [35]. The gene signatures corresponding to estrogen response– ER-early and ER-late (Hallmark pathway signatures from the Molecular Signature Database)– were downregulated, as expected, upon ER silencing. Importantly, ERa silencing led to a concurrent increase in activity of Hallmark-EMT, Hallmark-Glycolysis, Basal, EMT-up and Hallmark-Hypoxia gene sets, and a decrease in activity for luminal, FAO, Hallmark-ER-Early, Hallmark-ER-Late, EMT-down, and Hallmark-Oxphos gene sets as compared to WT cells. ERa-silenced cells also show increased expression of CD274, which codes for PD-L1 (Programmed death-ligand 1), a surface protein that plays a crucial role in cancer cells’ immune escape (Figure 3a and Figure S3a). This analysis suggests that the acquisition of tamoxifen resistance can drive changes along multiple axes of plasticity– EMT, metabolic plasticity, immune evasion, and a lineage switch from luminal to basal.

**Figure 3:**
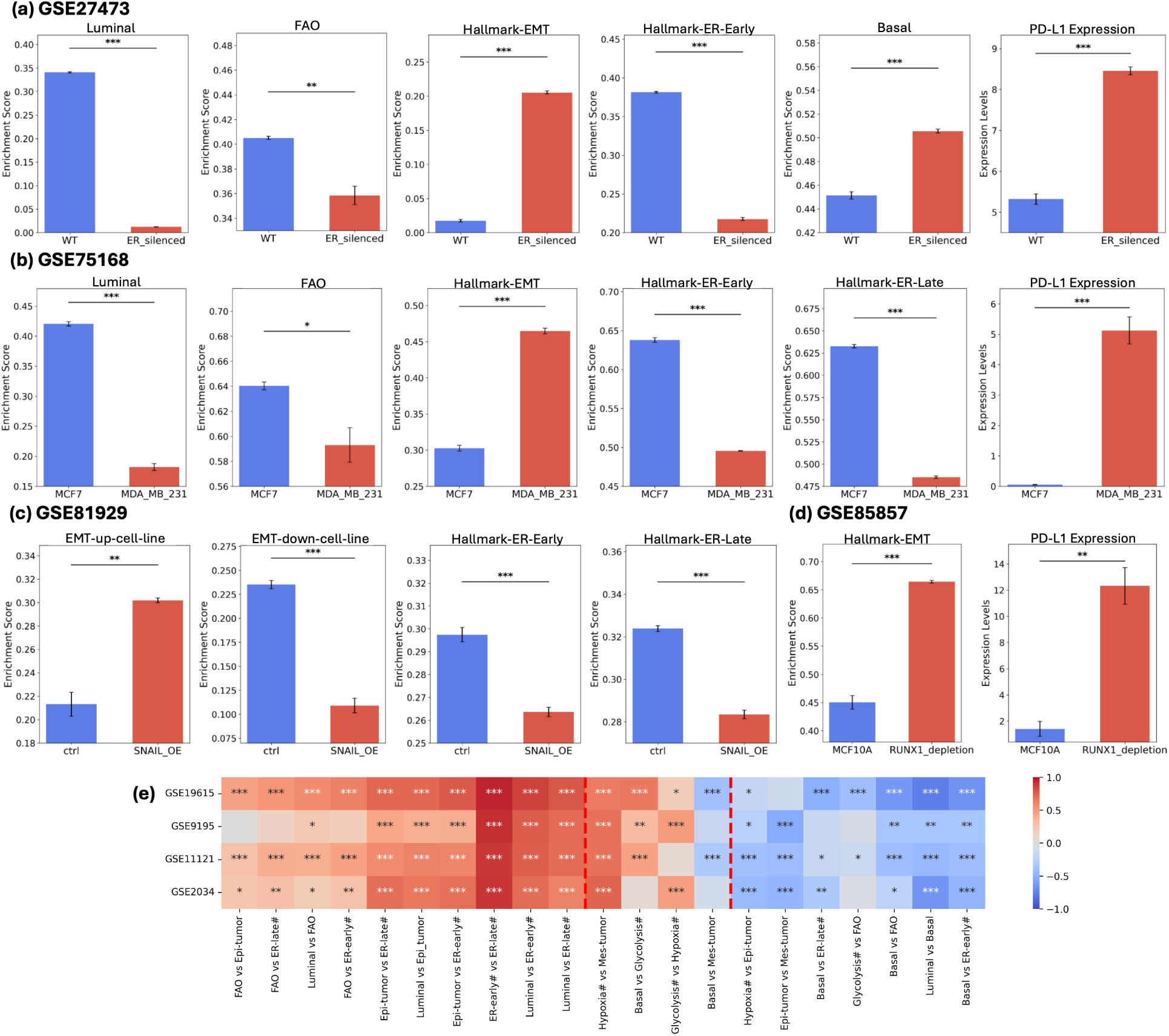
Analysis of bulk gene expression datasets showing associations between different axes of cellular plasticity. (a) Activity levels of Luminal, FAO, Hallmark-EMT, Hallmark-ER-Early and Basal gene sets; Expression levels of CD274 gene (PD-L1) in GSE27473 (MCF7). (b) Activity levels of Luminal, FAO, Hallmark-EMT, Hallmark-ER-Early and Hallmark-ER-Late gene sets; Expression levels of CD274 gene (PD-L1) in GSE75168 (MCF7 and MDA-MB-231). (c) Activity levels of EMT-up-cell-line, EMT-down-cell-line, Hallmark-ER-Early and Hallmark-ER-Late gene sets in GSE81929 (MCF10A). (d) Activity levels of Hallmark-EMT; Expression levels of CD274 gene (PD-L1) in GSE85857 (MCF10A). (e) Pairwise correlation analysis of Activity levels of gene sets in GSE19612, GSE9195, GSE11121 and GSE2034. Note: gene sets with ‘#’ in the end represent Hallmark gene sets obtained from MSigDB. (* indicating p < 0.05, ** indicating p < 0.01, *** indicating p < 0.001)

Triple Negative Breast Cancer (TNBC) lacks or shows very low levels of estrogen receptor (ER), Progesterone Receptor (PR) and Human Epidermal Growth Factor Receptor 2 (HER2). Thus, similar comparisons of normalised enrichment scores in GSE75168 [36] revealed that, the MDA-MB-231 (a TNBC cell line) showed increased activity of Hallmark-EMT, EMT-up, PD-L1-signature, Hallmark-Hypoxia and Basal gene sets, whereas they show decreased activity of Luminal, FAO, Hallmark-ER-Early, Hallmark-ER-Late and EMT-down gene-sets as compared to MCF7 (Figure 3b and Figure S3b). These observations were consistent with earlier work describing TNBC cells as being more basal-like, drug-resistant and highly Glycolytic [37, 38, 39]. Moreover, MDA-MB-231 also show increased PD-L1 (CD274) gene expression levels as compared to MCF7.

SNAIL (a pro-EMT transcription factor) is often found upregulated in cancer cells and is frequently used to induce EMT in epithelial cells [40]. Upon comparison of normalised enrichment scores for microarray dataset GSE81929 [41] (MCF10A cell line), it was revealed that EMT induction via overexpression of SNAIL results in an increase in activity of EMT-up gene-set, whereas a decrease in activity of EMT-down, Hallmark-ER-Early and Hallmark-ER-Late gene-sets (Figure 3c).

RUNX1 is known to control cell differentiation, stabilisation of epithelial phenotype and metastasis via acting as a transcriptional regulator of the IGF1R gene [42]. The highest levels of RUNX1 are observed in mammary epithelial cells, whereas RUNX1 levels are lower in tumorigenic and metastatic breast cancer cells. Comparison of normalised enrichment scores upon ssGSEA analysis of RNA-seq dataset GSE85857 [43] containing gene expression data for MCF10A cell line with and without RUNX1 depletion (Figure 3d and Figure S3c) revealed that RUNX1-depleted cells have higher activity for Hallmark-EMT, EMT-up, Hallmark-Hypoxia gene-sets, as well as exhibit increased expression levels of PD-L1(CD274) than WT MCF10A cells.

In another case study, we investigated cells with altered expression of GRHL2– Grainy head-like 2– a transcription factor known to suppress EMT. Also, higher expression of GRHL2 has been found to be associated with an epithelial-like phenotype [44, 45, 46, 47]. Analysis of gene expression dataset for GRHL2-knockdown cells and WT cells in ovarian cancer cell line (GSE118407) [48] demonstrated that shGRHL2 cells show increased activity for Hallmark-EMT and Hallmark-Hypoxia gene-sets and show decreased activity of Hallmark-Oxphos gene-set (Figure S3d).

Furthermore, we performed pairwise correlation analysis of normalised enrichment scores across four independent breast cancer patient datasets, namely: GSE19615 [49], GSE9195 [50], GSE11121[51] and GSE2034 [52]. This analysis revealed that Hallmark-Estrogen Response (ER-Early and ER-Late), Luminal, Fatty Acid Oxidation, and Epi-tumour signatures were strongly and positively correlated with one another. Similarly, Hallmark–Hypoxia, Mes-tumour, and Hallmark–Glycolysis signatures showed strong positive correlations with one another, except for Basal and Mes-tumour, which were negatively correlated. Consequently, these two groups of gene signatures were characterised by overall negative correlations between them.

These trends strengthen our observations from GRN simulations that the axes of phenotypic plasticity are strongly interconnected and can drive each other in specific directions corresponding to the characterisation of two ‘teams’ observed.

### 2.4 Single-cell data recapitulates systemic phenotypic plasticity patterns

After bulk gene expression datasets, we moved onto analysing single-cell breast cancer datasets. We performed GSEA on a single-cell breast cancer cell line dataset containing 32 cell lines (GSE173634) [53] on the following datasets: (), and obtained a pairwise Spearman correlation matrix between GSEA scores for different gene sets. Hierarchical clustering of this correlation matrix revealed the existence of two groups (Figure 4a(i)) one, containing Hallmark-Oxphos, FAO, Hallmark-ER-Early, Hallmark-ER-late, Luminal, EMT-down and KS-Epi-cell-line gene sets, while the other containing PD-L1-signature, BPMS, KS-Mes-cell-line, Hallmark-EMT, EMT-up-cell-line, Hallmark-Hypoxia, RPMS, Hallmark-Glycolysis, EMT-partial and Basal gene sets. Similar clustering of gene sets was observed for patient-derived tumour single-cell dataset GSE176078 (Figure 4b) [54].

**Figure 4:**
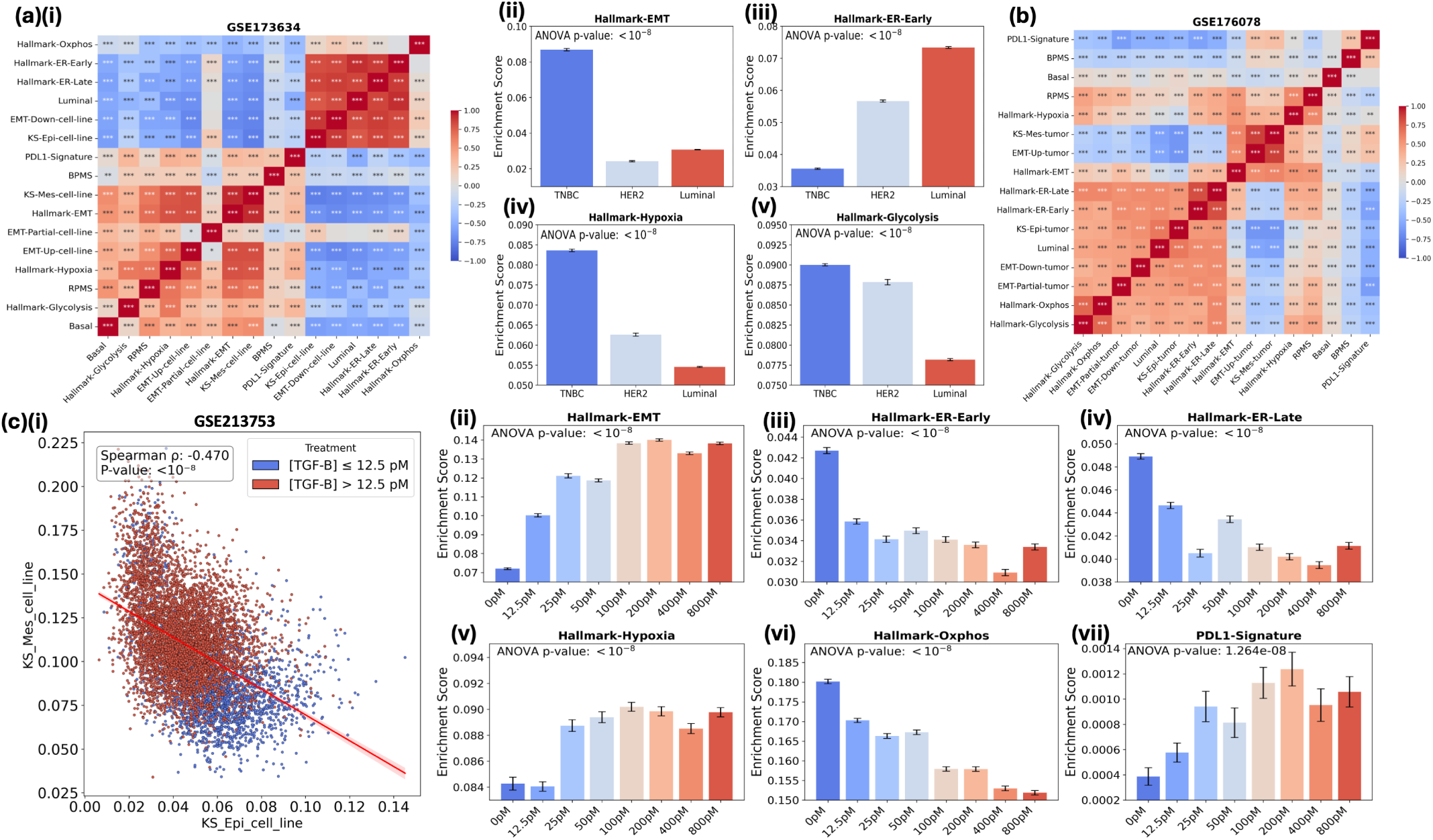
Single-cell analysis of breast cancer datasets linking different axes of cel-lular plasticity. (a) (i)Heatmap depicting pairwise Spearman’s correlation coefficients between different gene sets for GSE173634. Correlation plots for the association of cell line. (ii-v) Activ-ity of Hallmark-EMT, Hallmark-ER-Early, Hallmark-Hypoxia, Hallmark-Glycolysis. (b) Heatmap depicting pairwise Spearman’s correlation coefficients between different gene sets for GSE176078. (c) (i) Scatter plots showing correlations of ssGSEA scores of KS-Mes-cell-line and KS-Epi-cell-line in GSE213753. (ii-vii) Activity of Hallmark-EMT, Hallmark-ER-Early, Hallmark-ER-Late, Hallmark-Hypoxia, Hallmark-Oxphos and PD-L1 signature gene sets in GSE213753.

Biologically, this organisation grouped PD-L1, Hypoxia, Glycolysis, and Basal programs with those promoting EMT, mesenchymal, and hybrid phenotypes, while Oxphos, Fatty acid oxidation, Estrogen response, and Luminal programs clustered with epithelial and non-EMT states. These clustering patterns further reinforce the notion of coordination across multiple axes of cellular plasticity with EMT. Moreover, these results were in complete agreement with results obtained from analysis of in-silico simulations of GRN in earlier sections.

Further comparison of normalised enrichment scores (NES) for GSE173634 with cell lines separated into TNBC, HER2+ and ER+, revealed distinct pathway-specific activity patterns. TNBC cell lines exhibited higher activity of the Hallmark-EMT gene set compared with both HER2+ and ER+ cell lines (Figure 4a(ii)), while showing reduced activity of Hallmark-ER-Early and Hallmark-ER-Late gene sets (Figure 4a(iii), Figure S4b). In addition, TNBC cell lines demonstrated elevated activity of Hallmark-Hypoxia, Hallmark-Glycolysis, KS-Mes-cell-line, and PD-L1-signature compared with HER2+ and ER+ cell lines (Figure 4a(iv,v), and Figure S4c,e). Conversely, ER+ and HER2+ cell lines showed higher activity of KS-Epi-Cell-line and Hallmark-Oxphos gene sets relative to TNBC cell lines (Figure S4a,d).

TGF-*β* is a well-established inducer of epithelial-to-mesenchymal transition (EMT)[55]. It promotes loss of epithelial traits (such as E-cadherin) and acquisition of mesenchymal features (such as N-cadherin and vimentin) through SMAD-dependent and SMAD-independent signalling, which enhances cell migration, invasion, and plasticity. GSEA was performed on the single-cell RNA-seq dataset for MCF10A cell line treated with increasing concentrations of TGF-*β* (GSE213753) [56]. The analysis revealed a strong negative correlation between the activities of the KS-Epi-cell-line and KS-Mes-cell-line gene sets, reflecting the well-studied antagonism between epithelial and mesenchymal programs. This trend was evident in cells exposed to lower TGF-*β* concentrations compared to those treated with higher concentrations (Figure 4c(i)). Again, as expected, there was an increase in the activity of the Hallmark-EMT gene set with rising TGF-*β* concentrations (Figure 4c(ii)). In parallel, decreasing trends were observed for Hallmark-ER-Early, Hallmark-ER-Late, and Hallmark-Oxphos gene sets. Conversely, increasing trends were observed for Hallmark-Hypoxia and PD-L1-signature gene sets as TGF-*β* concentration increased (Figure 4c(iii-vii)). Together, these trends endorse our hypothesis that EMT progression usually co-occurs with decreased activity of ER signaling and oxidative phosphorylation, but with increased glycolytic activity [5].

### 2.5 Coordinated phenotypic transitions predict breast cancer aggressiveness and clinical outcomes

We next interrogated whether such a coordinated switch in cellular behaviour across multiple axes of plasticity associates with clinical responses. We first checked the association of individual axes of plasticity, focusing on the Estrogen Response (ER) pathway. The Hallmark Estrogen Response Early (Hallmark-ER-Early) and Hallmark Estrogen Response Late (Hallmark-ER-Late) gene sets from the Molecular Signatures Database (MSigDB) were used to classify patients based on median scores, for breast cancer patient cohorts from METABRIC and TCGA, and respective Kaplan-Meier plots were produced, reporting Hazard Ratios (HR) and corresponding p-values. Patients with high ER-Early or high ER-Late GSEA scores associated with better overall survival compared to those with low scores (HR= 0.82, p= 0.0027 for ER-Early and HR= 0.85, p= 0.015 for ER-Late gene sets) in the ER-positive cohort from METABRIC. This trend was also consistent in the entire METABRIC cohort (HR= 0.81, p= 0.00035 for ER-Early and HR= 0.88, p= 0.031 for ER-Late gene sets), in the ER-positive cohort in TCGA ( HR = 0.47, p= 0.00083 for ER-Early and HR= 0.33, p < 0.0001 for ER-Late gene sets) and in the entire cohort in TCGA (HR= 0.61, p= 0.0029 for ER-Early and HR= 0.59, p= 0.0014 for ER-Late gene sets) (Figure 5a, Figure S4a, Figure S6a(i, ii) and Figure S6c(i, ii)).

**Figure 5:**
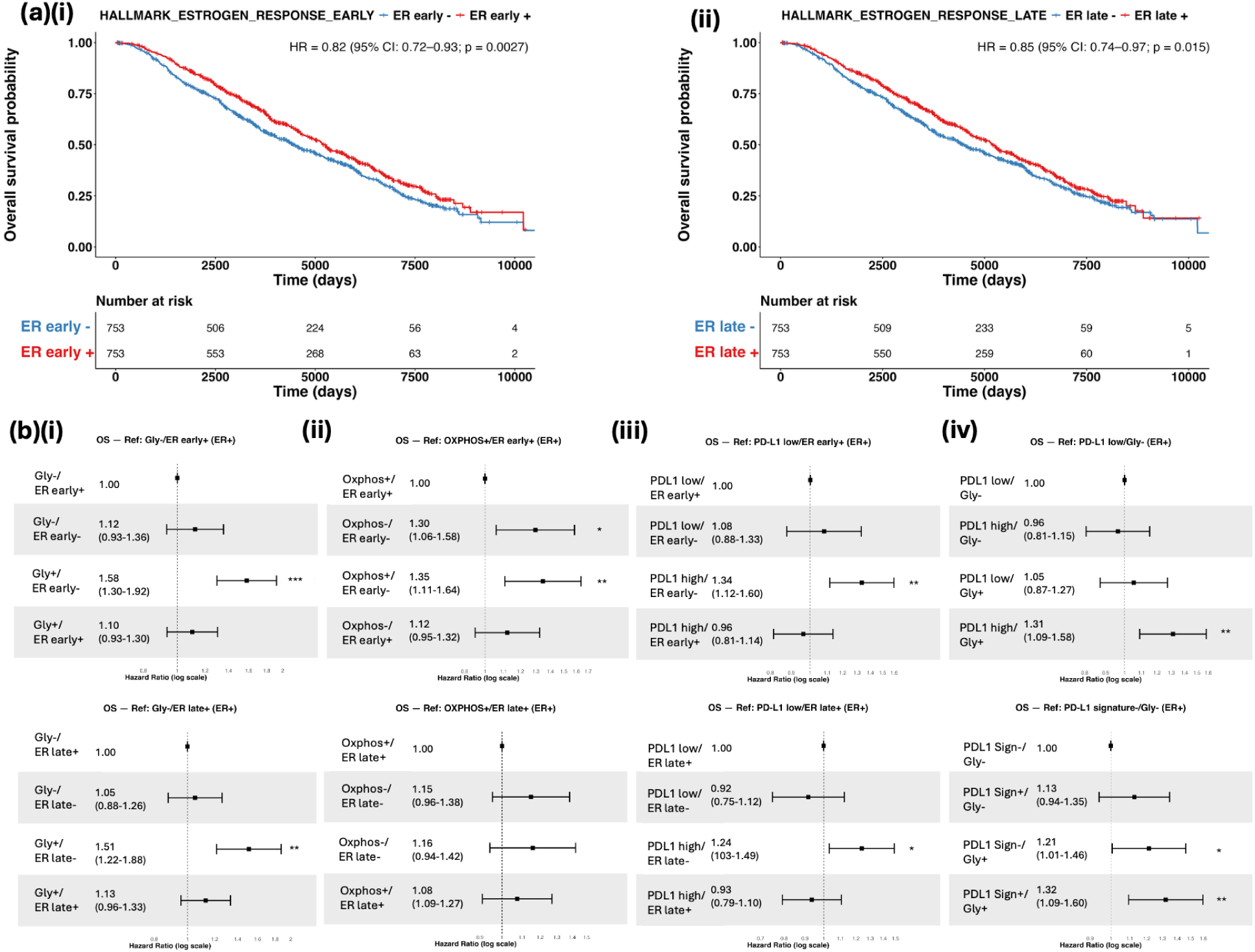
Survival analysis of estrogen receptor Positive (ER+) breast cancer patient data from METABRIC. (a) Kaplan-Meier overall survival (OS) curves of patients with breast cancer having either high or low activity of (i) Hallmark-ER-Early and (ii) Hallmark-ER-Late gene sets. (b) Forest plots comparing overall survival (OS) for different combinations of (i) Glycolysis vs Hallmark-ER-Early and Glycolysis vs Hallmark-ER-Late, (ii) Oxphos vs Hallmark-ER-Early and Oxphos vs Hallmark-ER-Late, (iii) PD-L1 expression vs Hallmark-ER-Early and PD-L1 expression vs Hallmark-ER-Late, and (iv) PD-L1 expression vs Glycolysis and PD-L1-(gene signature) vs Glycolysis.

Next, we investigated the association between different axes of cellular plasticity and overall patient survival (OS) using Cox regression analysis. Pairwise combinations of gene sets corresponding to different processes were tested, and hazard ratios were computed. A phenotype characterised by high-glycolysis together with low ER-Early or low ER-Late scores (Gly+/ER-Early– and Gly+/ER-Late–) was associated with significantly worse OS as compared to the reference with low-glycolysis and high ER-Early (or ER-Late) phenotype in both the ER-positive and the complete breast cancer cohorts of METABRIC (Figure 5b(i), Figure S5b(i)). The reference was chosen based on the composition of teams identified where ER*α*66– the ER isoform inhibited by tamoxifen [19]– shows antagonistic behavior with glycolysis, and previous literature showcasing association of tamoxifen resistance (driven by ER*α*36) with glycolysis [57, 58]. These trends were found to be consistent in the TCGA dataset as well, both for the full cohort and the ER-positive subset (Figures S6b(i) and S6d(i)). A similar association was observed for double combinations of Oxphos and ER-Early (or ER-Late) where the phenotype associated with low Oxphos and low ER-Early (or ER-Late) scores (Oxphos–/ER-Early– and Oxphos–/ER-Late–) correlated with worse OS relative to the reference with high Oxphos and high ER-Early (or ER-Late) phenotype. The same trends were observed in both METABRIC (Figure 5b(ii), Figure S4b(ii)) and TCGA (Figure S5b(ii) and Figure S5d(ii)) datasets across the entire cohorts and their corresponding ER-positive subsets. It is worth noting that in these combinatorial cases, coordinated switching along both axes of plasticity, but not along just one axis of plasticity, led to worse survival; for instance, in the context of low glycolysis & high ER-Early (or ER-Late) as the reference case, switching just the Glycolysis (from low to high) or ER-Early/ Late (from high to low) does not lead to statistically significant differences in the hazard ratio, but in cases where both the axes were flipped– glycolysis and ER-Early/ER-Late, the patient survival was worse (HR= 1.58 for for ER-Early and 1.51 for ER-Late) (Figure 5b (i)).

Next, we examined the association of PD-L1 expression with ER-Early and ER-Late pathways. As expected, phenotypes with high PD-L1 expression and low ER-Early or ER-Late scores were associated with worse survival compared to the reference phenotype of low PD-L1 expression and high ER-Early (or ER-Late) scores. These trends were consistent in both ER-positive and complete breast cancer cohorts of METABRIC (Figure 5b(iii) and Figure S4b(iii)) and were also recapitulated when using a PD-L1 gene signature [15] instead of PD-L1 expression levels in the complete cohort(Figure S4b(iv)). Again, in this combinatorial case, switching only the PD-L1 levels (from low to high) or ER-Early/ER-Late scores (from high to low) does not lead to statistically significant differences in the hazard ratio, but in cases when both PD-L1 and ER-Early (or ER-Late) were flipped, the patients associated with worse survival (HR= 1.34, for ER-Early and HR= 1.24, for ER-Late) (Figure 5b (iii)).

Lastly, the combined impact of PD-L1 expression and Glycolysis score on overall survival was also assessed. Phenotypes with high PD-L1 expression (or high PD-L1 signature) together with high Glycolysis scores were associated with significantly worse OS compared to the reference phenotype with low PD-L1 expression (or signature) and low Glycolysis scores. We remark that the reference case was selected based on our teams-level analysis showcasing PD-L1 and glycolysis belonging to same team, and our previous results reporting a positive correlation of PD-L1 levels with glycolysis scores [16, 59]. These associations were observed in both ER-positive and complete breast cancer cohorts of METABRIC (Figure 5b(iv) and Figure S4b(v)). Similar to the previous combinatorial analyses of two axes of plasticity, this case further highlights the more aggressive behavior resulting from the coordinated alteration of multiple axes of plasticity—specifically, the flipping of both PD-L1 expression (or PD-L1 signature) and glycolysis —rather than changes in either axis alone, was linked to worse clinical outcomes.

Motivated by these observed associations of two axes of plasticity together (Oxphos and Glycolysis with Hallmark-ER-Early and Hallmark-ER-Late), we probed whether triple combinations of Oxphos, Glycolysis, and ER-Early (or ER-Late) also exhibited concordant trends. As before, Cox regression analysis was performed using the samples classified as high or low along each of the three axes (thus, a total of 8 cases). In the ER-Early case, three phenotypes: low Oxphos/high Glycolysis/low ER-Early, high Oxphos/high Glycolysis/low ER-Early, and low Oxphos/high Glycolysis/high ER-Early were associated with significantly worse OS relative to the reference phenotype (high Oxphos/low Glycolysis/high ER-Early) (Figure 6a(i)). Importantly, analogous results were obtained when analysing ER-Late gene set instead of ER-Early with combinations low Oxphos/high Glycolysis/low ER-Late, high Oxphos/high Glycolysis/low ER-Late, and low Oxphos/high Glycolysis/high ER-Late exhibiting worse survival(Figure 6a(ii)).

**Figure 6:**
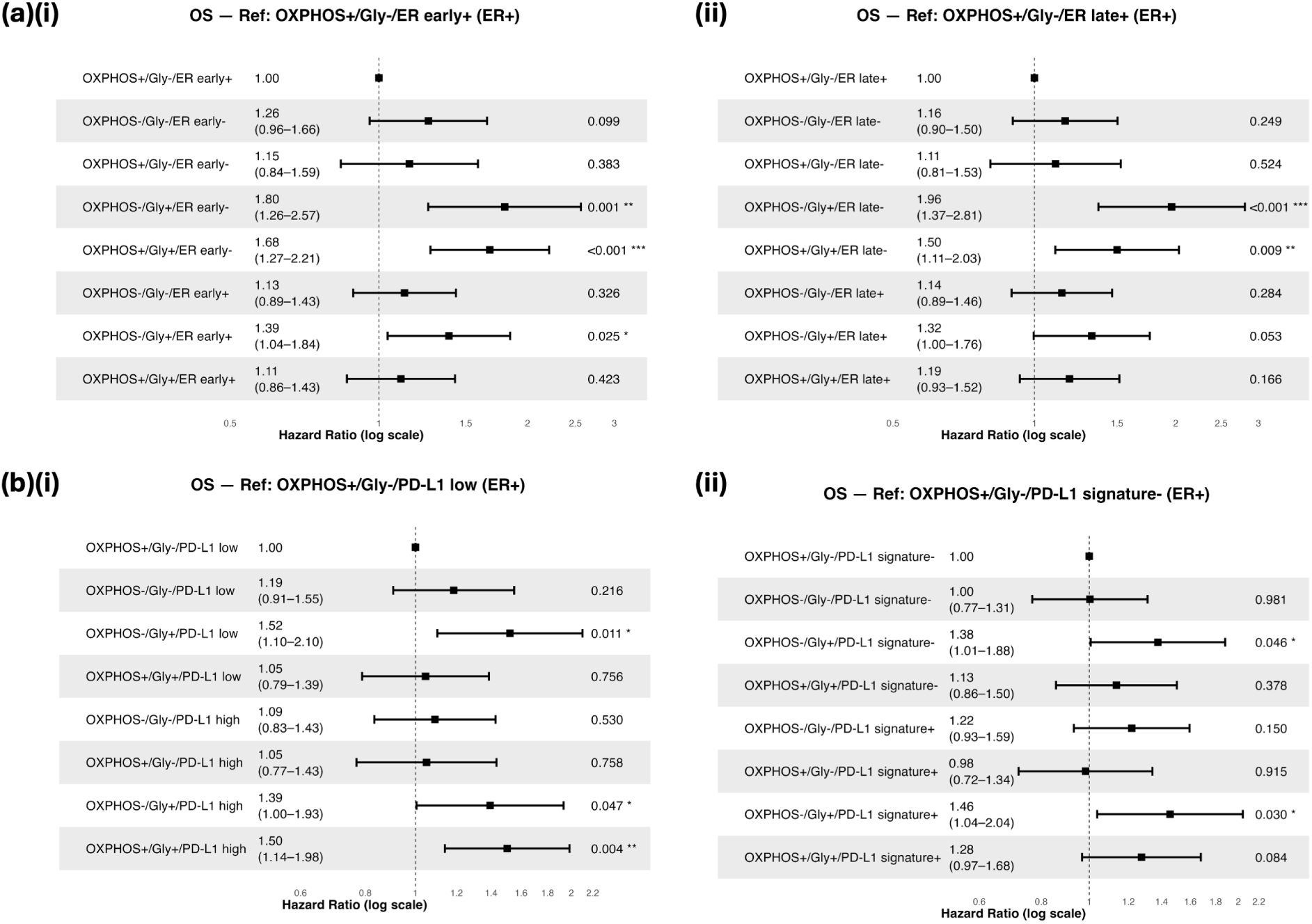
Triple combination survival analysis for METABRIC estrogen receptor pos-itive (ER+) breast cancer patient data. (a) Forest plots comparing overall survival (OS) for different combinations of (i) Oxphos vs Glycolysis vs Hallmark-ER-Early and (ii) Oxphos vs Glycolysis vs Hallmark-ER-Late. (b) Forest plots comparing overall survival (OS) for different combinations of (i) Oxphos vs Glycolysis vs PD-L1-gene and (ii) Oxphos vs Glycolysis vs PD-L1-signature.

In the above combinatorial cases, the phenotypes associated with worse survival all represent conditions with high Glycolysis gene set activity, accompanied by reduced activity of at least one of the Oxphos or ER-Early (or ER-Late) gene sets. Notably, high Glycolysis consistently drove worse survival outcomes in most Oxphos/ER-Early permutations, except when both Oxphos and ER-Early were simultaneously high (high Oxphos/high Glycolysis/high ER-Early) where Oxphos and ER-Early perhaps exhibited a stronger reinforcement of the ‘team’ associated with improved overall survival. These trends were observed in both the full METABRIC cohort (Figure S7a) and the ER-positive cohort (Figure 6a).

Further, we performed triple-combination Cox regression analyses of Oxphos and Glycolysis with PD-L1 gene expression and the PD-L1 gene signature. In case of PD-L1 expression, the combinations of low Oxphos/high Glycolysis/low PD-L1, low Oxphos/high Glycolysis/high PD-L1, and high Oxphos/high Glycolysis/high PD-L1 were associated with worse survival compared to the reference (high Oxphos/low Glycolysis/low PD-L1). Similar trends were observed when PD-L1 gene signature was used. Combinations of low Oxphos/high Glycolysis/low PD-L1-signature, low Oxphos/high Glycolysis/high PD-L1-signature, and high Oxphos/high Glycolysis/high PD-L1-signature were linked to worse survival relative to the reference (high Oxphos/low Glycolysis/low PD-L1-signature)(Figure 6b). These observations were again observed in the full METABRIC cohort (Figure S7b).

Next, to check whether these trends could be observed in other cancer types, we conducted a pan-cancer overall survival (OS) analysis in TCGA data. Double combinations of Hallmark-EMT vs Hallmark-Glycolysis, Hallmark-EMT vs Hallmark-Glycolysis vs Hallmark-Hypoxia, and Hallmark-Glycolysis vs Hallmark-Oxphos were evaluated. The reference phenotype was defined as having low EMT, low Glycolysis, low Hypoxia, and high Oxphos for each respective combination. Across all four pairwise comparisons, consistent trends were observed: phenotypes with high-EMT, high-glycolysis, high-hypoxia, and low-Oxphos were associated with significantly worse OS relative to the reference groups, where some cancers show these patterns with significant statistical power, whereas others did not. Particularly, cancers BLCA, CESC, HNSC, KIRP, LGG, LIHC, LUAD, MESO, and PAAD emerged as cancers which show these patterns with varying statistical power across different combinations. Some cancer types showed these patterns with strong statistical significance, while others did not. Specifically, BLCA, CESC, HNSC, KIRP, LIHC, LUAD, MESO, and PAAD were identified as cancers showing these associations with statistical significance in all four cases. (Figure S8 (a, b, c, d))

Motivated by these findings, GSEA analysis was conducted for the 30 cancer types in the TCGA cohort and pairwise correlation matrices were constructed. Interestingly, upon clustering of these correlation matrices it was observed that BLCA, CESC, HNSC, KIRP, LAUD, PAAD and MESO with an exception of LIHC showing clustering of gene sets associated with epithelial programmes clustered together and gene sets associated with mesenchymal programmes clustered together (Figure S9). These trends highlight that cancer types where a teams-like coordinated behavior was more predominantly observed also led to worse clinical outcomes in combinatorial cases compared to the individual axes of plasticity.

## 3 Discussion

Our study highlights that multiple axes of cancer cell plasticity — namely metabolic reprogramming, epithelial-to-mesenchymal plasticity (EMP), luminal-basal phenotype switching, and stemness, do not function independently but instead act in an interconnected and coordinated fashion. The key nodes responsible for driving plasticity along each individual axis– such as ZEB1 driving EMP and HIF1a driving metabolic plasticity– form two “teams”, with intra-team interactions being predominantly activating, while inter-team interactions being predominantly inhibitory, directly and/or indirectly. These opposing teams collectively shape the aggressiveness of cancer progression, with one set of phenotypic states (high Glycolysis, stem-like, basal-like, drug-resistant, and mesenchymal/hybrid) associated with more aggressive disease, whereas the other (high oxidative phosphorylation [Oxphos], non–stem-like, luminal-like, drug-sensitive, and epithelial) linked to less aggressive outcomes. Perturbation of one axis was found to influence cellular phenotypes across multiple axes, underscoring the systemic nature of phenotypic plasticity.

Our results can coherently synthesize many experimental observations made in breast cancer: a) TGFb signaling, a key inducer of EMT, drives Glycolysis [60], b) cells in hybrid E/M state can have heightened stemness/tumor-initiation potential compared to those in epithelial state [61, 27], c) tamoxifen-resistant cells exhibit characteristics of EMP [62], and d) luminal-to-basal lineage switch manifesting markers associated with a hybrid E/M state [63]. While not necessarily universal, similar interconnections have been reported in other cancer types as well, such as coupling between EMT-metabolism in carcinomas [59], the interlinked dynamics of PD-L1 levels with proliferative-invasive transition in melanoma [64], and the involvement of EMP associated genes in neuroendocrine transition in small cell lung cancer [65]. Therefore, this higher-order coordination of phenotypic behavior in cancer cells can be a more generic feature emergent from the underlying topology of context-specific GRNs.

This ‘teams’ like behavior in GRNs not only reduces the dimensionality of consequent phenotypic space [65], but also provides a systematic framework to investigate synthetic lethality of cancer cells, for instance, identifying therapeutic vulnerabilities of EMT+ and drug-resistant prostate cancer [66]. Mapping the modules of co-expressed genes enabled by ‘teams’ can accelerate finding viable synthetic lethality combinations, enabling the goals of precision oncology. Our clinical data analysis shows that coordinated switching of cellular behavior, such as a simultaneous upregulation of glycolysis and downregulation of ER pathways, lead to worse clinical outcomes than either of them being modulated. This observation points to the existence of ‘teams’ facilitating a higher ‘fitness’ of tumor cells, particularly in cancer metastasis that has extraordinary attrition rates. Whether these ‘teams’ identified from a bottom-up approach connect with archetypes observed from a top-down approach [67], remains to be investigated further.

Our work has several limitations. First, we only focus on steady-state behavior of these GRNs without delving into time-course dynamics, while there may be dynamic regulation of different glycolytic reactions during early and late stages of EMT [68], or maximum tumor-initiation potential observed for intermediate stages of EMT [69]. Second, we have not incorporated the epigenetic layer of regulation of these processes, such as the interplay between hypoxia, EMT and chromatin modifier CTCF [70, 71]. Third, our analysis has been restricted to cell-autonomous behavior, without explicitly considering cell-cell communication signaling among tumor cells or with other stromal cells. Preliminary evidence suggests that the concept of ‘teams’ can be extended to investigate tumor-immune interactions as well [71].

These limitations notwithstanding, our findings provide a unifying systems-level organizational principle connecting multiple forms of plasticity to tumor adaptability and resistance. Our results unravel that cancer cell fitness can be enhanced by coordinated transitions across phenotypic plasticity dimensions, thereby indicating novel targets to disrupt a systems-level cell reprogramming.

## 4 Methods

### 4.1 RACIPE

To simulate dynamical behaviour exhibited by a particular Gene regulatory network (GRN), we applied the RACIPE framework for GRN random simulation [32], Fig 1A. RACIPE samples parameters over a biologically relevant range to generate multiple parameter sets and all possible steady-state behaviors of the network were quantified across the ensemble of parameter sets. The simulation results were analysed to understand the emergent states arising from the network topology and to determine their respective frequencies within the simulation cohort.

The input network topology file used for the simulations contained information about inhibitory and activating links between nodes. For each kinetic model generated from the input network, RACIPE samples multiple initial parameters from the designated range for each parameter. The expression levels of each node in a GRN were determined by a set of ordinary differential equations given by:

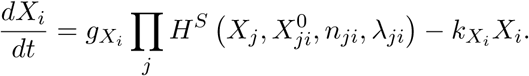

Here, *X_i_* denoted the concentration of the gene product corresponding to the node *X_i_* in the GRN, and *dX_i_/dt* represented the rate of change of gene expression over time. *g_X__i_* indicated the basal production rate, while *k_X__i_* denoted the basal degradation rate of gene product *X_i_*. The term *H^S^* referred to the shifted Hill function, which modelled the activation or inhibition of the gene node *X_i_* by other nodes in the network. *n_ji_* represented the Hill coefficient, and *X*^0^ was the threshold value for the Hill function. *X_j_* denoted the concentration of gene product *X_j_*, where *X_j_* was a node that either activated or inhibited *X_i_*.

A total of 10,000 parameter sets were used for simulations conducted in this study. Initial conditions for each node were randomly sampled from a log-uniform distribution spanning the minimum to maximum expression levels for that node. RACIPE produced log_2_-normalized steady-state gene expression levels for each gene product as output. This output data was subsequently *z*-normalized and used for downstream analysis.

### 4.2 Network Topology Analysis

#### Adjacency Matrix

The adjacency matrix (Adj) was used to represent the network topology and the regulatory interactions among nodes in the gene regulatory network of interest. In this matrix, rows represent the source nodes for regulatory links, while columns represent the corresponding target nodes.

A direct inhibitory interaction was represented by *−*1, while a direct activating interaction was represented by +1, and the absence of a direct interaction was represented by 0.

#### Influence Matrix

The adjacency matrix accounts for only the direct interactions between two nodes in the network. Hence, any analysis done on the adjacency matrix inherently lacks information regarding indirect interactions between two nodes. To overcome this shortfall, another formulation called the *Influence Matrix* (Infl) was used. The Influence Matrix is a representation similar to the adjacency matrix, with rows representing source nodes and columns representing target nodes [72]. The Influence Matrix was calculated for a defined path length *l*_max_ by summing the matrices 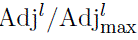 for *l ≤ l*_max_ as:

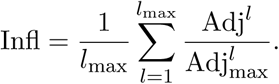

Here, 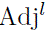 represents the adjacency matrix (Adj) raised to the power *l*. Adj_max_ represents the adjacency matrix with all inhibition links replaced by activation links. The ratio 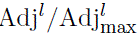 represents the element-wise division of values in 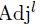 by values in 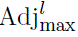. The summation is divided by *l*_max_ to restrict the elements of the final Influence Matrix between *−*1 and 1.

#### Identification of Teams from the Influence Matrix

“Teams” were defined as a group of genes that exhibit coordinated regulatory behaviour [72], i.e., supporting members within their own team and suppressing members of the opposing team. This can be interpreted from two perspectives:

##### Source Node perspective (How a Node Affects Others)

- Nodes positively regulated members of their own team.
- Nodes negatively regulated members of the opposing team.
- Nodes inhibited their own team members with less strength than they inhibited members of the opposing team.
- Nodes activated their own team members more strongly than they activated those of the opposing team.

##### Target Node perspective (How a Node is Affected by Others)

- Nodes were positively regulated by members of their own team.
- Nodes were negatively regulated by members of the opposing team.
- Inhibitory input from own team members was weaker compared to that from the opposing team.
- Activating input from own team members was stronger than that from the opposing team.

To identify the “Teams” from the Influence Matrix, an *n ×* 2*n* matrix was constructed from the original *n × n* Influence Matrix *I*. This was done by horizontally concatenating *I* with its transpose *I^T^*:

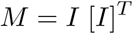

This new matrix *M* captured both outgoing and incoming influences for each node in its rows. Hierarchical clustering was then performed on the *n* rows of matrix *M*, resulting in the division of genes into two distinct clusters. These clusters were referred to as the two “Teams”, each comprising genes with corresponding regulatory roles.

### 4.3 Generation of Random Networks

To analyse the features and characteristics that the unique architecture of the WT biological networks give rise to, a set of random, non-biological networks that do not exist in nature, referred to as “Random networks,” were generated. The following steps were employed to generate a random network:

1. Start with the biological network as a template.
2. All regulatory interactions (edges) were shuffled among themselves using the random function in Python, such that the total number of inhibition and activation links remains the same.
3. The shuffled activation and inhibition links were then reassigned to the network. Reassigning of the edges was done such that node pairs that lacked regulatory connections prior to shuffling remained unconnected afterward as well.
4. The newly generated random network was subjected to a uniqueness check: it was compared against an existing list of previously generated random networks. If the network was not already present in the list, it was added. If it was found to be a duplicate, it was discarded, and steps 2 through 4 were repeated to generate a new unique network.

This process was repeated *n* times to obtain a total of *n* distinct random networks.

### 4.4 Gene Expression Data Analysis

TCGA gene expression data of 33 different cancer types from the UCSC-Xena browser [73]. Publicly available gene expression datasets were obtained from NCBI GEO website [74]. Different preprocessing steps were carried for RNA-seq and Microarray datasets. For RNA-seq datasets the respective counts matrices were obtained from NCBI GEO and converted to TPM which were log transformed and used for downstream analyses. For microarray datasets the respective raw “.CEL.gz” files were downloaded from NCBI GEO and Robust Multiarray Average (RMA) normalisation was performed to obtain normalised intensity data matrix with rows as genes and columns as samples.

#### ssGSEA

Single-sample GSEA (ssGSEA), an extension of Gene Set Enrichment Analysis (GSEA) [75], calculates separate enrichment scores for each pairing of a sample and a gene set. Each ssGSEA enrichment score represents the degree to which the genes in a particular gene set are coordinately up– or down-regulated within a sample.

For bulk RNA-seq and microarray datasets, GSEApy [76], a Python package, was used to perform ssGSEA. For single-cell RNA-seq datasets, AUCell [77], an R package, was used to perform ssGSEA.

### 4.5 Survival Analysis

We analyzed two breast-cancer cohorts: TCGA-BRCA (downloaded from the UCSC Xena “TCGA Breast Cancer” hub) and METBARIC. We profiled the following signatures: Glycolysis, Oxidative phosphorylation, Estrogen response early, Estrogen response late, PD-L1 signature, PD-L1 gene (CD274). All gene signatures are listed in the Supplementary Table X.

For signature scoring, we performed ssGSEA using the GSVA [78], a R package. For each signature, samples were classified into “+” (high) and “-” low groups based on the cohort-specific median of the ssGSEA score. For PD-L1 gene expression (CD274), samples were split into high/low categories by the cohort-specific median expression level, without ssGSEA.

For Kaplan-Meier analyses, we implemented three levels of grouping models. Groups were defined as follows: one-axis (single signature), two-axis (pairwise combinations), and three-axis (triple combinations). The KM survival curves were generated using the ggsurvplot function in survminer package [79] in R. Cox proportional-hazard models were fitted using the survival package with reference groups defined a priori. Forest plots of hazard ratios and 95% confidence intervals were generated using the survminer package.

## Code Availability

The GSE datasets analysed were obtained directly from GEO (Gene Expression Omnibus). The codes used are available from the GitHub repository: https://github.com/i-donno-riteshkm/multiaxis_paper

## Authors’ Contribution

MKJ and JTG designed and conceived research and obtained funding. RKM, YF, SR and AJP performed research. RKM, YF, SR, AJP, AK, YS and YuS analyzed and interpreted data. RKM prepared a first draft of the manuscript. RKM, YJ, MKJ and JTG edited the draft with inputs from all the authors. All authors have read and approved the final manuscript.

## 5. Supplementary Figures

**Supplementary Figure 1:**
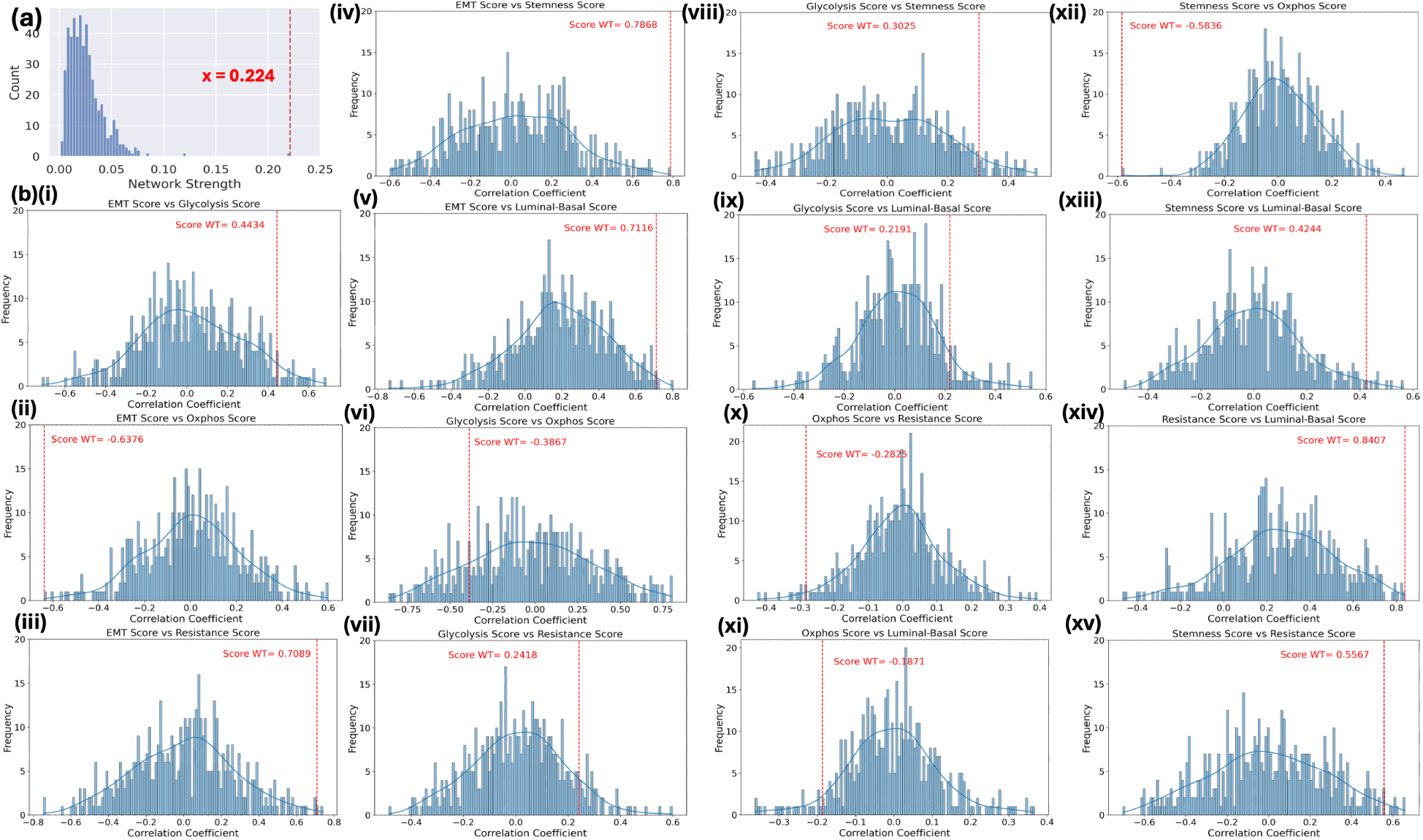
(a) Team-strength frequency histogram, with dashed red line representing Team strength of WT network. (b) Explicit Histograms showing correlation strength of WT network w.r.t. 500 other random networks with respect every pairwise combination of 6 Scores shown in Figure 2b(ii).

**Supplementary Figure 2:**
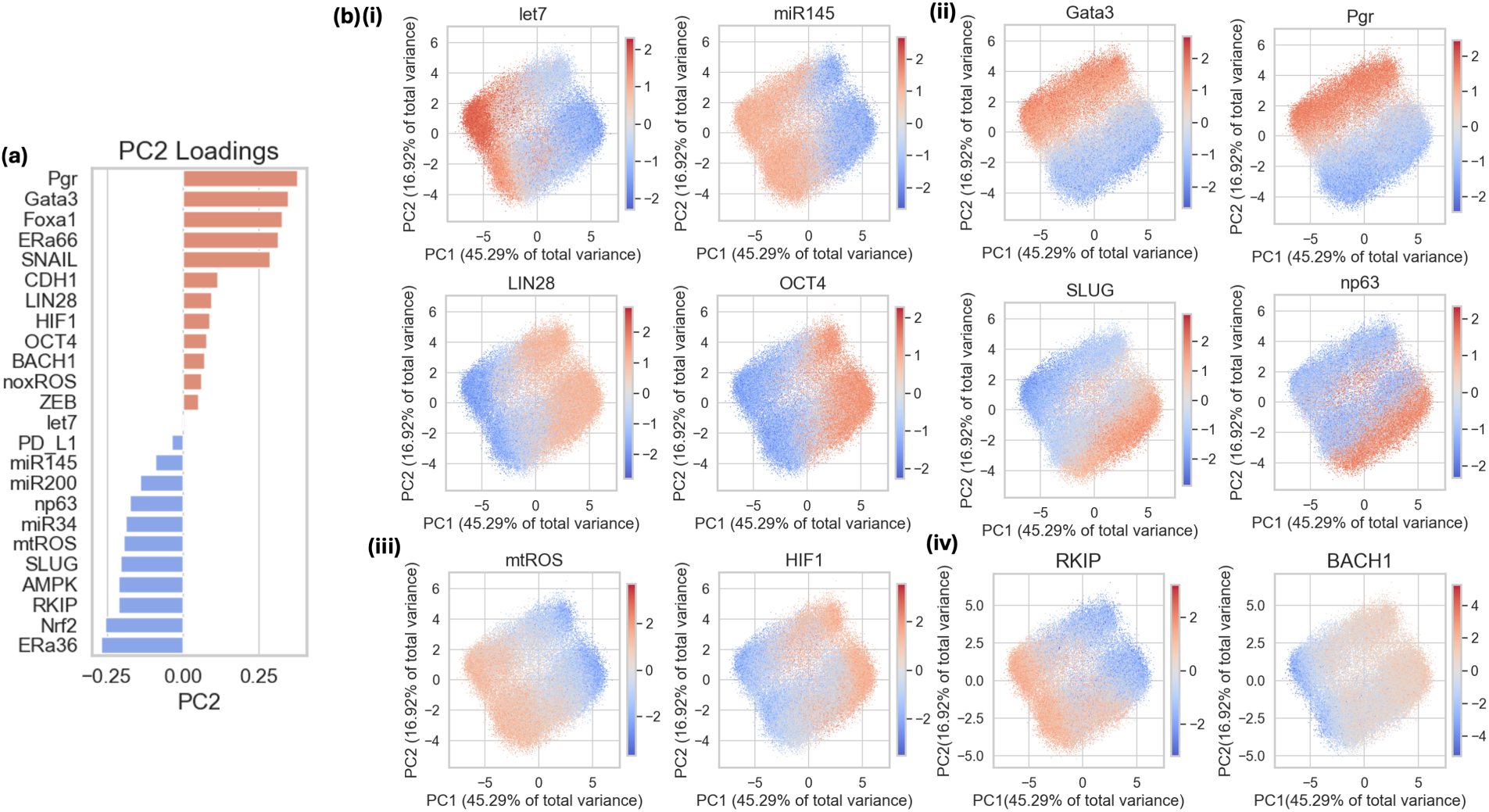
Results form Principle Component analysis for RACIPE Simulation results. (a) Vertical Bar plot showing the PC2 loading coefficients of different genes in the network. (b) Scatterplots of PC1 (capturing 45.29% variance) vs PC2 (capturing 16.92% variance) shaded by z-normalised score values for genes associated with different modules (i) Stemness Module (ii) Luminal-Basal Plasticity Module (iii) Metabolic Reprogramming Module (iv) RKIP and BACH1.

**Supplementary Figure 3:**
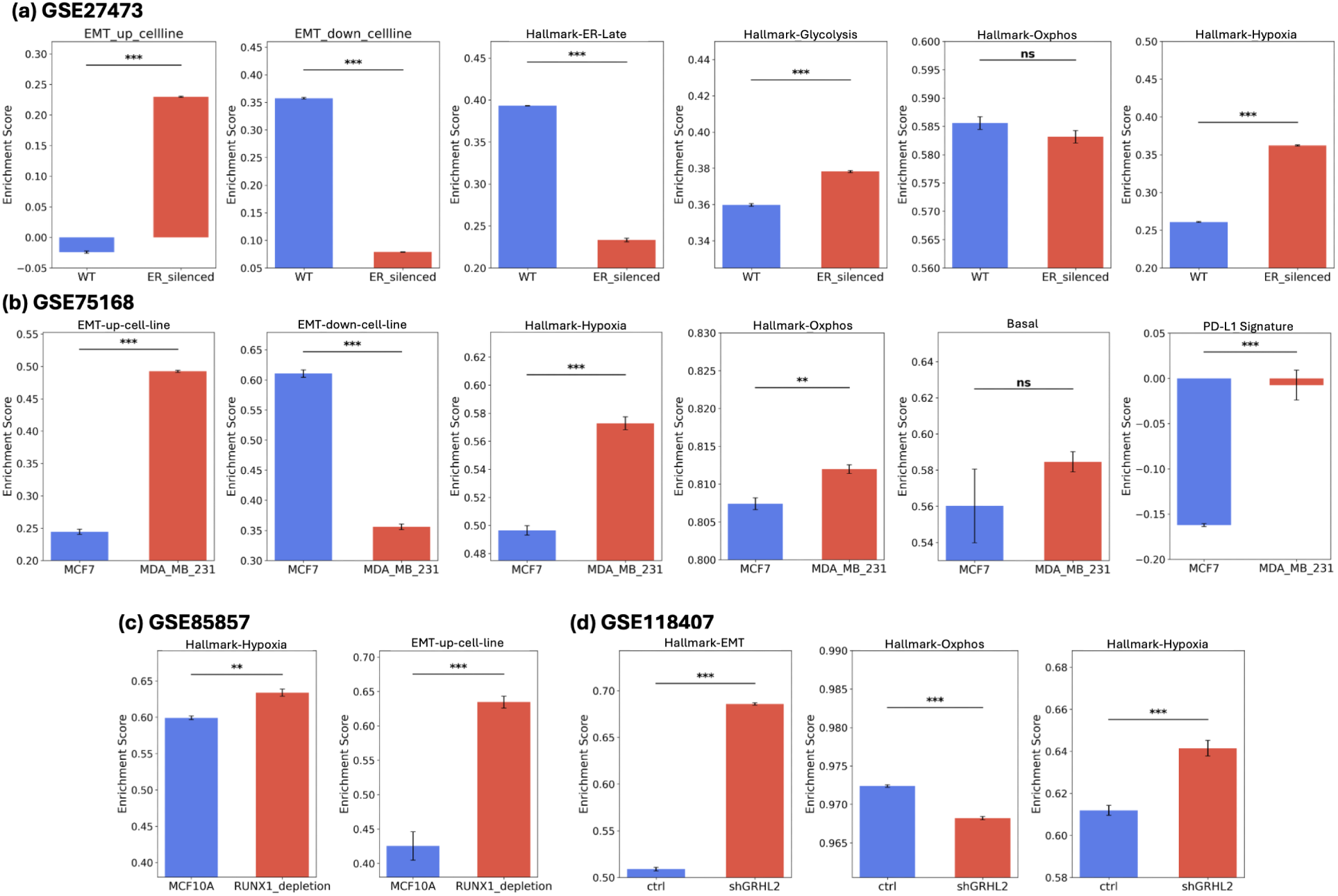
(a) Activity levels of EMT-Up-cell-line, EMT-Down-cell-line, Estrogen-Response-Late, Hallmark-Glycolysis, Hallmark-Oxphos and Hallmark-Hypoxia gene sets in GSE27473 (MCF7). (b) Activity levels of EMT-Up-cell-line, EMT-Down-cell-line, Hallmark-Hypoxia, Hallmark-Oxphos, Basal and PD-L1-Signature gene sets in GSE75168 (MCF7 and MDA-MB-231). (c) Activity levels of Hallmark-Hypoxia, EMT-Up-cell-line in GSE85857 (MCF10A). (d) Activity levels of Hallmark-EMT, Hallmark-Oxphos and Hallmark-Hypoxia gene sets in GSE118407 (OVAC).

**Supplementary Figure 4:**
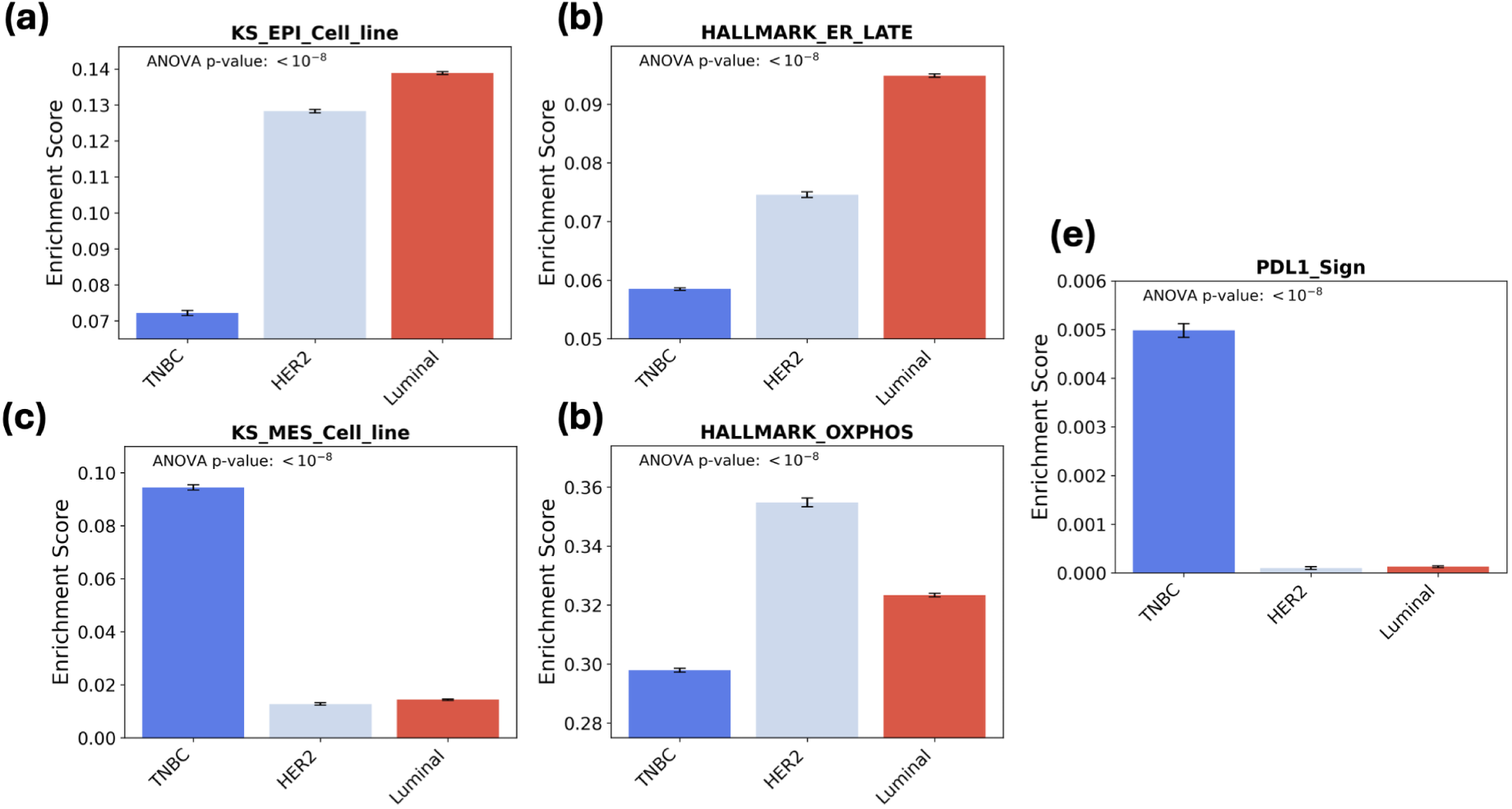
Comparison of normalised enrichment score for GSEA analysis of GDE173634 for (a) KS-Epi-cell-line, (b) Hallmark-ER-Late, (c) KS-Mes-cell-line, (d) Hallmark-Oxphos, (e) PD-L1-Signature.

**Supplementary Figure 5:**
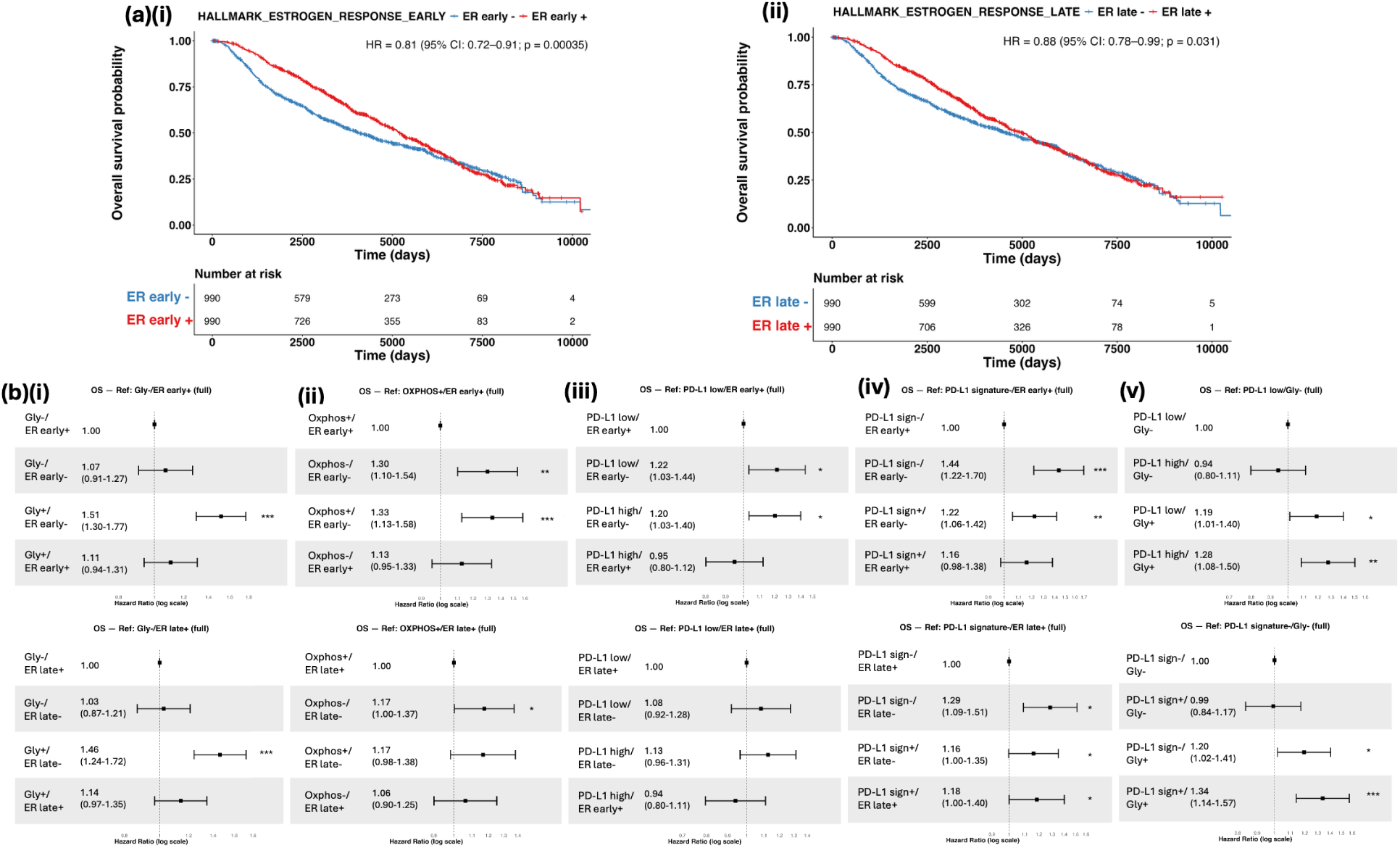
Survival analysis for the complete Breast cancer cohort from METABRIC. (a) Kaplan-Meier overall survival (OS) curves of patients with breast cancer and either high or low activity of (i) Hallmark-ER-Early and (ii) Hallmark-ER-Late gene sets. (b) Forest plots comparing overall survival (OS) for different combinations of (i) Glycolysis vs Hallmark-ER-Early and Glycolysis vs Hallmark-ER-Late (ii) PD-L1(gene expression) vs Hallmark-ER-Early and PD-L1(gene expression) vs Hallmark-ER-Late (iii) PD-L1(gene signature) vs Hallmark-ER-Early and PD-L1(gene signature) vs Hallmark-ER-Late (iv) PD-L1(gene expression) vs Glycolysis and PD-L1-(gene signature) vs Glycolysis (v) Oxphos vs Hallmark-ER-Early and Oxphos vs Hallmark-ER-Late.

**Supplementary Figure 6:**
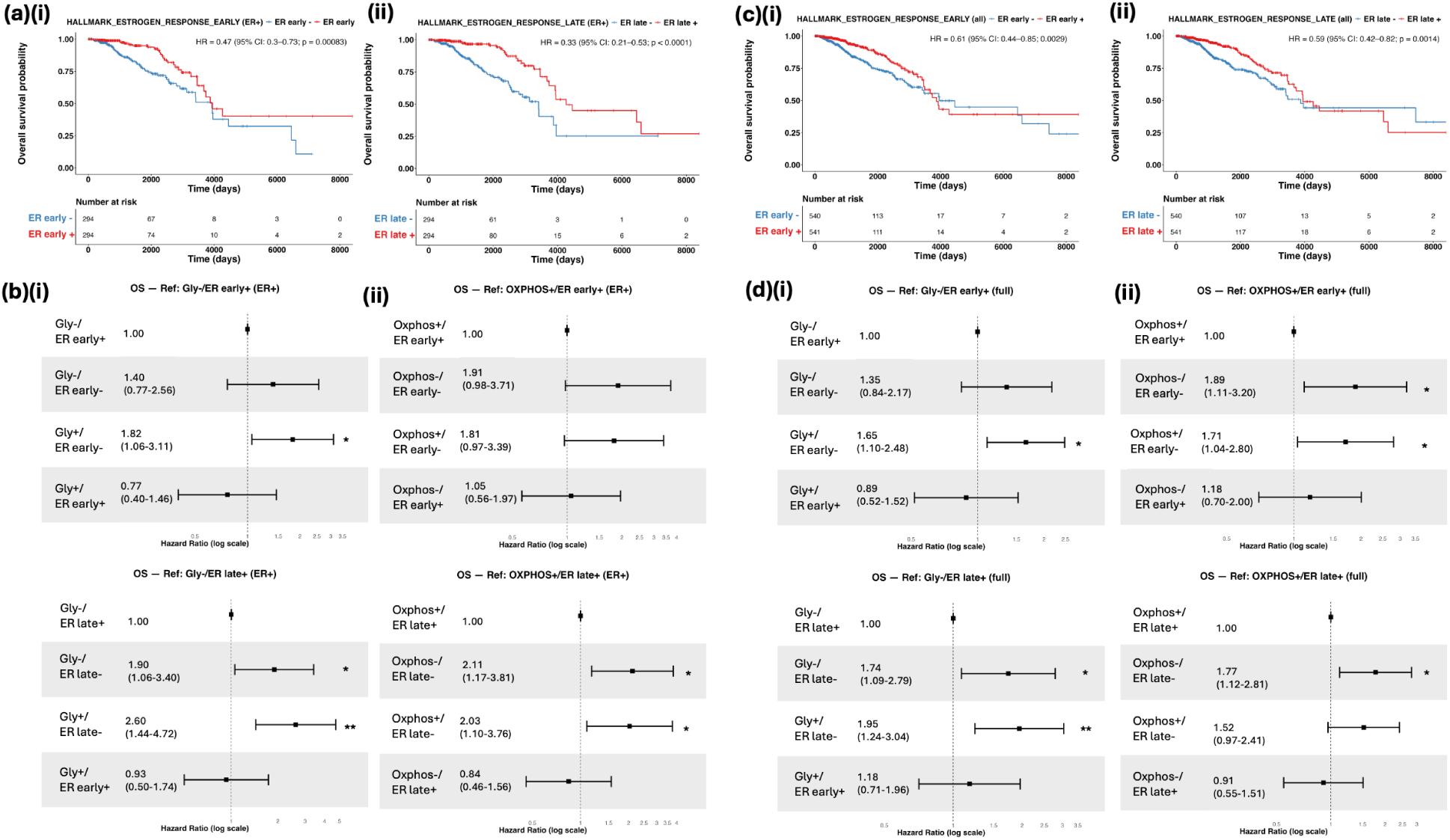
Survival analysis Breast cancer Patient data from TCGA Estrogen-Receptor-Positive (ER+) samples and all samples. (a) Kaplan-Meier overall survival (OS) curves of patients with ER+ breast cancer samples with either high or low activity of (i) Hallmark-ER-Early and (ii)Hallmark-ER-Late gene sets. (b) Forest plots comparing overall survival (OS) for different combinations of (i) Glycolysis vs Hallmark-ER-Early and Glycolysis vs Hallmark-ER-Late (ii) Oxphos vs Hallmark-ER-Early and Oxphos vs Hallmark-ER-Late for ER+ Breast Cancer samples. (c) Kaplan-Meier overall survival (OS) curves of patients with all breast cancer samples with either high or low activity of (i) Hallmark-ER-Early and (ii)Hallmark-ER-Late gene sets. (d) Forest plots comparing overall survival (OS) for different combinations of (i) Glycolysis vs Hallmark-ER-Early and Glycolysis vs Hallmark-ER-Late (ii) Oxphos vs Hallmark-ER-Early and Oxphos vs Hallmark-ER-Late for all Breast Cancer Samples.

**Supplementary Figure 7:**
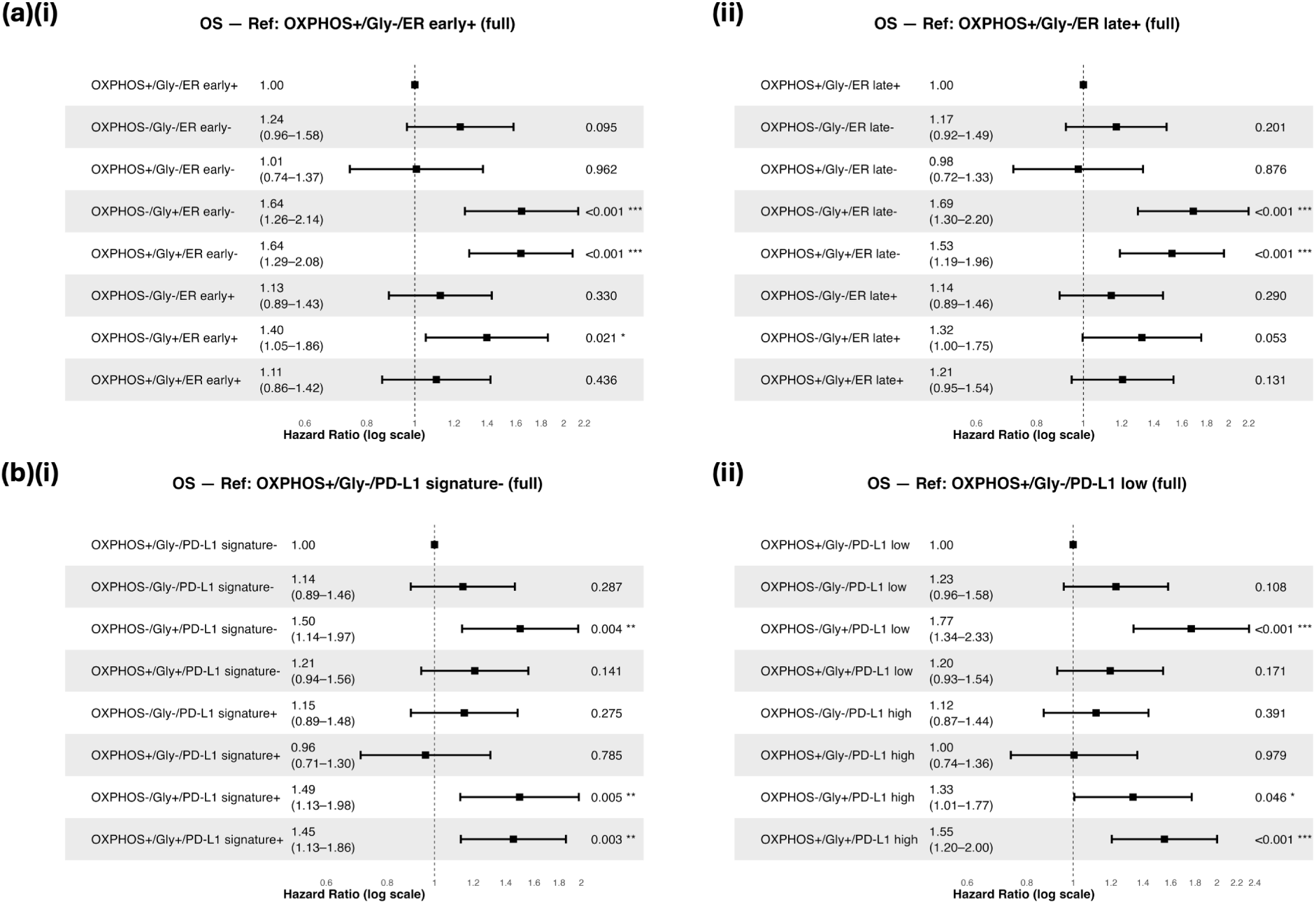
Survival analysis with complete Breast cancer cohort samples from METABRIC. (a) Forest plots comparing overall survival (OS) for different combinations of (i) Oxphos vs Glycolysis vs Hallmark-ER-Early and (ii) Oxphos vs Glycolysis vs Hallmark-ER-Late. (b) Forest plots comparing overall survival (OS) for different combinations of (i) Oxphos vs Glycolysis vs PD-L1-gene and (ii) Oxphos vs Glycolysis vs PD-L1-signature.

**Supplementary Figure 8:**
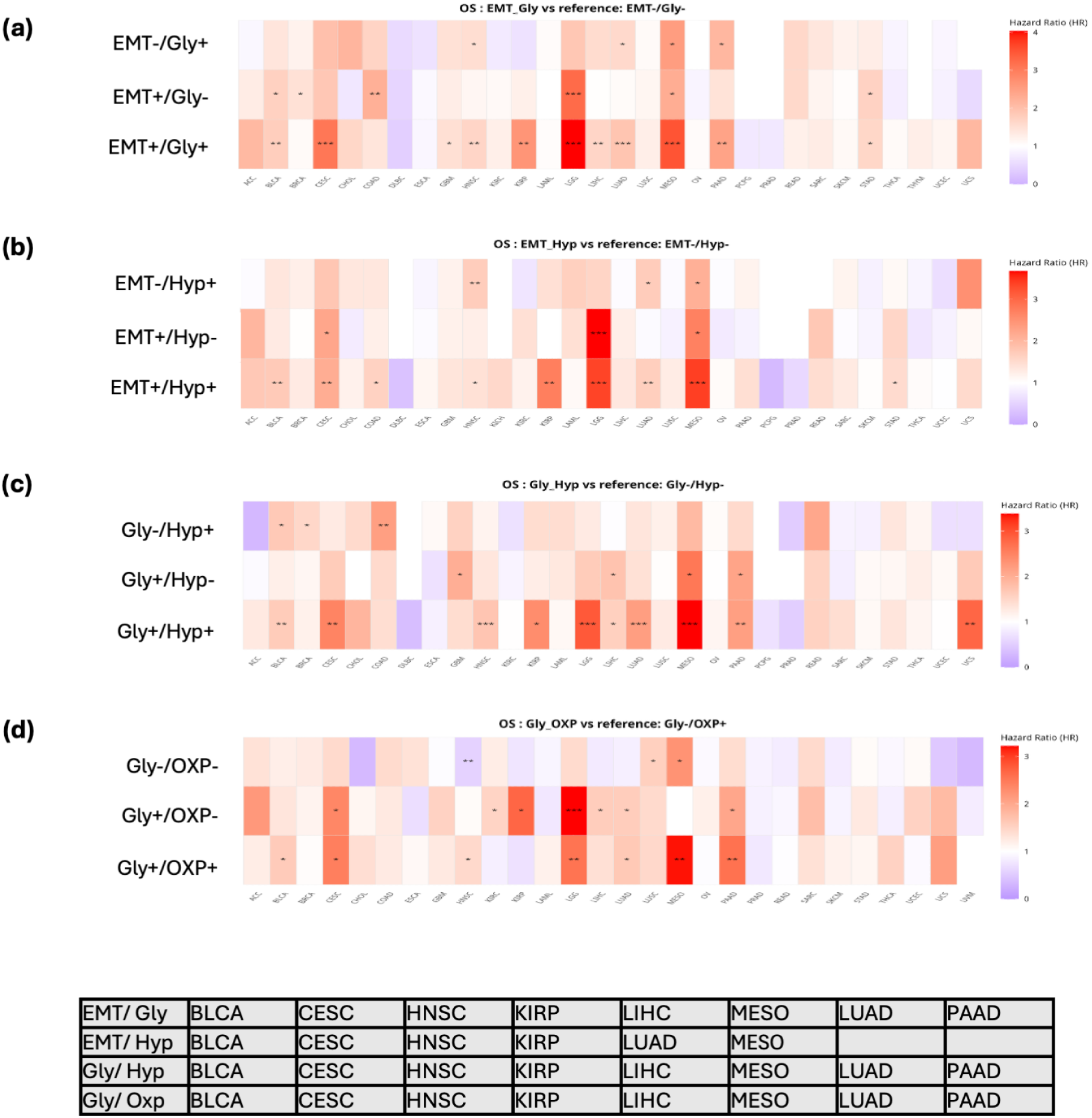
survival analysis heatmaps for different cancers in TCGA cohort showing Hazard ratio for different combinations of (a) Hallmark-EMT vs Hallmark-Glycolysis, (b) Hallmark-EMT vs Hallmark-Hypoxia, (c) Hallmark-Glycolysis vs Hallmark-Hypoxia and (d) Hallmark-Glycolysis vs Hallmark-Oxphos.

**Supplementary Figure 9:**
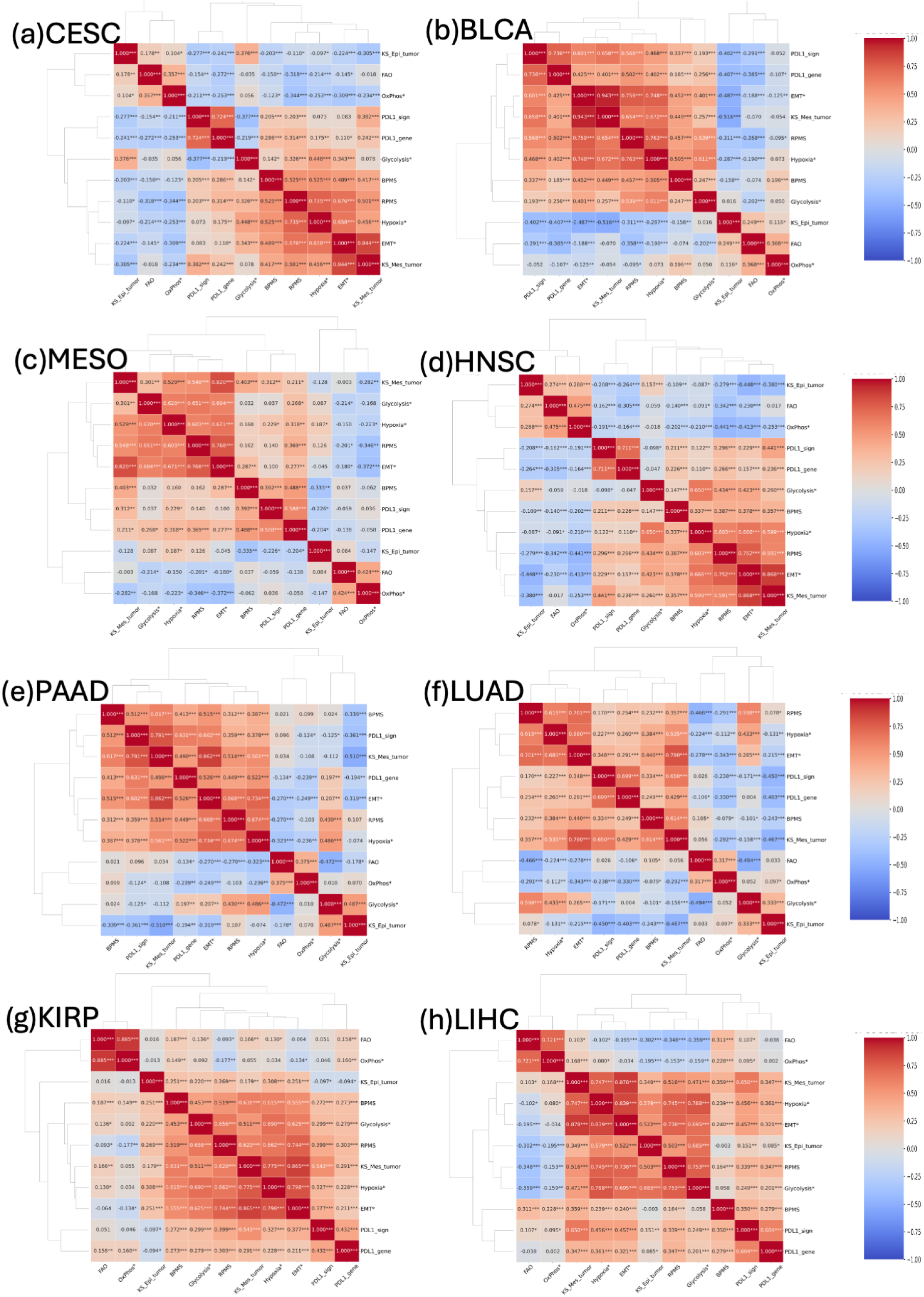
GSEA scores pairwise correlation heatmaps for different cancers in TCGA cohort (a) CESC, (b) BLCA, (c) MESO, (d) HNSC, (e) PAAD, (f) LUAD, (g) KIRP, (h) LIHC.

