## Supplementary Tables 1 and 2 for "Interconnected axes of phenotypic plasticity drive coordinated cellular behaviour and worse clinical outcomes in breast cancer"

### Supplementary table 1:

| GEO id | Type | Desription |
|---|---|---|
| GSE27473 | microarray | Expression data from MCF7 cell line before and after silencing of Estrogen recepto |
| GSE75168 | RNA-seq | RNA-Seq of different breast cancer cell lines MCF10A, MCF7 and MDA-MB-231. |
| GSE85857 | RNA-seq | Wt MCF10A cell-line and MCF10A cells with depletion of Runx1 |
| GSE118407 | RNA-seq | RNA of OVCA4209 control and GRHL2 knockdown (shGRHL2#12) cells were analyzed |
| GSE19615 | microarray | 115 patient primary human breast tumors sample |
| GSE9195 | microarray | 77 estrogen receptor-positive breast cancer patinets samples treated with tamoxifen |
| GSE11121 | microarray | 200 node-negative breast cancer patient samples |
| GSE2034 | microarray | 180 lymph-node negative relapse free BC patients and 106 lymph-node negate patients that developed a distant metastasis |
| GSE81929 | microarray | SNAI1 overexpression effect on MCF-10A mammary epithelial cell line (mRNA) |
| GSE173634 | Single-cell | Single-cell sequencing of 32 breast cancer cell-lines |
| GSE176078 | Single-cell | single-cell data from human breast cancers patients |
| GSE213753 | Single-cell | Dosage effect of TGF-beta on EMT induction of MCF10A cells |

### Supplementary Table 2: Links in the Network

| **Source** | **Target** | **Type** | **Reference** |
|---|---|---|---|
| OCT4 | miR145 | Inhibition | [1] |
| OCT4 | OCT4 | Activation | [2] |
| AMPK | BACH1 | Inhibition | [3] |
| AMPK | AMPK | Inhibition | [4] |
| AMPK | HIF1 | Inhibition | [4] |
| AMPK | miR200 | Activation | [4] |
| AMPK | mtROS | Activation | [4] |
| AMPK | mtROS | Inhibition | [4] |
| AMPK | noxROS | Inhibition | [4] |
| AMPK | SNAIL | Inhibition | [4] |
| AMPK | ZEB | Inhibition | [4] |
| BACH1 | BACH1 | Inhibition | [5] |
| BACH1 | RKIP | Inhibition | [5] |
| BACH1 | SLUG | Activation | [6] |
| CDH1 | ZEB | Inhibition | [7] |
| CDH1 | Nrf2 | Inhibition | [8] |
| ERa36 | ZEB | Activation | [9] |
| ERa36 | ERa66 | Inhibition | [10] |
| ERa66 | ERa36 | Inhibition | [9] |
| ERa66 | ERa66 | Activation | [9] |

| **Source** | **Target** | **Type** | **Reference** |
|---|---|---|---|
| ERa66 | SLUG | Inhibition | [9] |
| ERa66 | let7 | Activation | [11] |
| ERa66 | Gata3 | Activation | [12] |
| ERa66 | Pgr | Activation | [13] |
| ERa66 | PD-L1 | Inhibition | [7] |
| Foxa1 | CDH1 | Activation | [14] |
| Foxa1 | ERa66 | Activation | [15] |
| Foxa1 | SLUG | Inhibition | [16] |
| Gata3 | Foxa1 | Activation | [17] |
| Gata3 | ERa66 | Activation | [12] |
| Gata3 | CDH1 | Activation | [18] |
| Gata3 | Pgr | Activation | [19] |
| Gata3 | ZEB | Inhibition | [20] |
| Gata3 | ERa36 | Inhibition | [10] |
| HIF1 | SLUG | Activation | [21] |
| HIF1 | ZEB | Activation | [22] |
| HIF1 | AMPK | Inhibition | [4] |
| HIF1 | HIF1 | Activation | [4] |
| HIF1 | miR200 | Inhibition | [4] |
| HIF1 | mtROS | Inhibition | [4] |
| HIF1 | noxROS | Activation | [4] |
| HIF1 | SNAIL | Activation | [4] |
| HIF1 | ERa66 | Inhibition | [23] |
| HIF1 | PD-L1 | Activation | [24] |
| let7 | BACH1 | Inhibition | [5] |
| let7 | let7 | Activation | [25] |
| let7 | LIN28 | Inhibition | [25] |
| let7 | ZEB | Inhibition | [25] |
| let7 | PD-L1 | Inhibition | [26] |
| let7 | HIF1 | Inhibition | [27] |
| let7 | ERa36 | Inhibition | [28] |
| LIN28 | OCT4 | Activation | [29] |
| LIN28 | let7 | Inhibition | [25] |
| LIN28 | LIN28 | Activation | [25] |
| miR145 | OCT4 | Inhibition | [1] |
| miR145 | ZEB | Inhibition | [1] |
| miR145 | ERa66 | Inhibition | [30] |
| miR145 | PD-L1 | Inhibition | [31] |

| **Source** | **Target** | **Type** | **Reference** |
|---|---|---|---|
| miR200 | SLUG | Inhibition | [9] |
| miR200 | ZEB | Inhibition | [9] |
| miR200 | Nrf2 | Activation | [8] |
| miR200 | PD-L1 | Inhibition | [7] |
| miR200 | LIN28 | Inhibition | [28] |
| miR200 | HIF1 | Inhibition | [4] |
| miR34 | mtROS | Activation | [4] |
| miR34 | noxROS | Activation | [4] |
| miR34 | SNAIL | Inhibition | [4] |
| mtROS | AMPK | Activation | [4] |
| mtROS | HIF1 | Activation | [4] |
| noxROS | AMPK | Activation | [4] |
| noxROS | HIF1 | Activation | [4] |
| np63 | SLUG | Activation | [32] |
| np63 | ZEB | Inhibition | [32] |
| np63 | np63 | Activation | [8] |
| Nrf2 | ZEB | Inhibition | [8] |
| Nrf2 | Pgr | Inhibition | [33] |
| Nrf2 | PD-L1 | Activation | [34] |
| Nrf2 | SNAIL | Inhibition | [25] |
| Nrf2 | HIF1 | Activation | [35] |
| PD-L1 | CDH1 | Inhibition | [7] |
| PD-L1 | OCT4 | Activation | [36] |
| PD-L1 | SNAIL | Activation | [37] |
| Pgr | ERa66 | Activation | [38] |
| Pgr | Pgr | Activation | [13] |
| RKIP | SNAIL | Inhibition | [39] |
| RKIP | let7 | Activation | [5] |
| SLUG | CDH1 | Inhibition | [7] |
| SLUG | ERa66 | Inhibition | [9] |
| SLUG | miR200 | Inhibition | [9] |
| SLUG | SLUG | Activation | [9] |
| SLUG | ZEB | Activation | [9] |
| SLUG | SNAIL | Inhibition | [40] |
| SLUG | np63 | Activation | [32] |

| **Source** | **Target** | **Type** | **Reference** |
|---|---|---|---|
| SNAIL | RKIP | Inhibition | [41] |
| SNAIL | SLUG | Inhibition | [40] |
| SNAIL | miR200 | Inhibition | [25] |
| SNAIL | SNAIL | Inhibition | [25] |
| SNAIL | ZEB | Activation | [25] |
| SNAIL | miR34 | Inhibition | [4] |
| ZEB | CDH1 | Inhibition | [7] |
| ZEB | ERa66 | Inhibition | [9] |
| ZEB | miR200 | Inhibition | [9] |
| ZEB | ZEB | Activation | [9] |
| ZEB | miR145 | Inhibition | [29] |
| ZEB | miR34 | Inhibition | [4] |
